# An image-based framework for in silico trials of ablation strategies in scar-related ventricular tachycardia

**DOI:** 10.64898/2026.08.06.743176

**Authors:** Niccolò Biasi, Matteo Parollo, Davide M. Vultaggio, Giulio Zucchelli, Alessandro Tognetti

## Abstract

Scar-related ventricular tachycardia (VT) is sustained by patient-specific structural and functional remodeling of the ventricular substrate, and the optimal substrate-based ablation strategy remains debated. We present an image-based computational framework for conducting controlled in silico trials of VT ablation strategies. Patient-specific left ventricular electrophysiology models were generated from late gadolinium enhancement cardiac magnetic resonance images by incorporating image-derived scar, border-zone tissue with structural fibrosis, fiber orientation, and physiologically plausible Purkinje-driven sinus activation. A dedicated standalone graphical user interface was developed to perform interactive virtual ablation based on imaging-derived or simulated electrophysiological data. We implemented standardized VT reinducibility testing to compare different lesion sets in terms of residual VT inducibility and ablation burden. As a proof of concept, the framework was applied to 20 patients with ischemic or non-ischemic cardiomyopathy undergoing VT ablation. Sustained VT was inducible in 17 patients, yielding 127 sustained VT episodes and 88 unique reentrant circuits at baseline. Four substrate-based ablation strategies were compared: scar homogenization, primary deceleration-zone ablation, primary plus secondary deceleration-zone ablation, and CMR-guided scar dechanneling. All strategies significantly reduced VT inducibility compared with baseline. Scar homogenization achieved the largest reduction in residual unique sustained VTs but required the largest ablated myocardial volume. Conversely, CMR-guided scar dechanneling showed the most favorable efficiency profile by reducing VT inducibility while limiting ablated viable myocardium. The proposed framework enables quantitative comparison of ablation efficacy, ablation burden, and mechanisms of ablation success or failure in image-guided VT therapy planning.

## 1. Introduction

Structural heart disease (SHD) refers to a group of pathological conditions that involve structural abnormalities, mainly resulting in myocardial fibrosis. SHD may be a consequence of ischemic (i.e., myocardial infarction) or non-ischemic mechanisms such as myocarditis, dilated, hypertrophic, or arrhythmogenic cardiomyopathy. The European Society of Cardiology has recently estimated that in 2019, ischemic heart disease alone caused 34 million potential years of life lost, accounting for 57% of all cardiovascular diseases [1]. Myocardial fibrosis as the result of SHD is established as a critical substrate for cardiac arrhythmogenesis and sudden cardiac death. Indeed, the presence of inexcitable scar tissue combined with functional and structural remodeling in the border zone favors the initiation and maintenance of reentrant circuits [2]. It is estimated that at least 90% of sudden cardiac deaths occur in individuals with underlying SHD, and most commonly ischemic heart disease (50% to 80% of sudden cardiac deaths) [3]. Despite advancements in pharmacological therapy, recurrent ventricular tachycardia (VT) in SHD patients remains frequent and is associated with increased morbidity, mortality, and impaired quality of life. Catheter ablation has emerged as a cornerstone treatment for scar-related VT in patients affected by SHD, both as a first-line therapy [4] or as a second-line option when pharmacological treatments are ineffective or undesired [2]. Catheter ablation aims to eliminate the arrhythmogenic substrate by destroying small areas of remnant myocardial tissue to interrupt potential reentry circuits. The standard clinical approach to VT ablation involves arrhythmia induction and activation mapping to delineate the reentrant circuit and identify the critical isthmus serving as the ablation target. On the other hand, substrate-based approaches use high-density electroanatomic mapping during sinus or paced rhythm to identify potential ablation targets such as delayed or abnormal electrograms [5] or areas with slow conduction, such as deceleration zones (DZs) [6]. Substrate-based approaches avoid important drawbacks associated with arrhythmia induction, such as the hemodynamic compromise, risk of degeneration into ventricular fibrillation (VF), and the need for electrical cardioversion [7]. In the context of substrate-based approaches, imaging modalities for scar identification have gained attention for their potential to guide or aid VT ablation. Imaging-based VT ablation strategies use imaging to detect potential reentry circuits based only on anatomical information. The ventricular substrate can be inferred either directly through late gadolinium enhancement cardiac magnetic resonance (LGE-CMR) [8, 9] or indirectly by analyzing myocardial wall thickness via computed tomography [10]. By relying only on anatomical information, imaging-based approaches circumvent the need for high-density mapping employing multipolar catheters, which is often time-intensive and can increase procedural arrhythmogenicity [6]. At the same time, single-center or observational studies have shown that LGE-CMR-guided or aided approaches are both feasible and safe, with lower VT recurrence rates than the standard clinical approach [11, 12, 9].

Cardiac electrophysiology computational modeling has emerged as an innovative tool for supporting risk stratification, procedural planning, and therapy guidance [13, 14]. Regarding SHD, and particularly ischemic heart disease, most of the previous research efforts focused on the identification of optimal ablation sites [15, 16]. Waight et al. [17] have recently reported that electrogram abnormalities and regions of conduction slowing are more frequent in the optimal ablation sites predicted by computer models. The Virtual Induction and Treatment of Arrhythmia (VITA) framework was developed in [18] to identify potential sites of block and assess their vulnerability to reentry. The framework has been shown to accurately predict VT recurrence [19] in post-infarction patients. The Arrhythmic Risk Score (ARRISK) was recently proposed by Serra et al. [20] as a simulation-based VT risk assessment and demonstrated strong concordance with clinical outcomes in a 51-patient dataset. The ARRISK score can be considered as an extension of the Virtual-heart Arrhythmia Risk Predictor (VARP) protocol, originally proposed by Arevalo et al. [21]. Computer cardiac electrophysiology models can also serve as a research tool for investigating the pathophysiological mechanisms underlying arrhythmia generation. Villar-Valero et al. [22] used image-based computational models to examine the role of scar anatomy and border zone properties in the induction of VTs. Their study highlights that structural remodeling (i.e., fibrosis) plays a major role in the development of arrhythmias, whereas ionic remodeling is less critical. Finally, in silico clinical trials have been shown to provide important insights across different medical areas [23], including cardiac electrophysiology. For example, Dasì et al. have recently carried out an in silico clinical trial to determine optimal pharmacological and ablation therapies in atrial fibrillation patients [24]. Similarly, cardiac digital twins have been used to evaluate how amiodarone influences VT inducibility and bipolar electrogram markers [25].

This work aims to develop an image-based framework for conducting in silico trials of substrate-based ablation strategies in SHD by exploiting computational modeling and image analysis. Starting from clinical imaging data, the framework builds cardiac electrophysiology models with personalized arrhythmic substrate, including scar and fibrotic tissue. A dedicated graphical user interface (GUI) was developed for supporting interactive virtual ablation by clinical staff in the generated cardiac models. This GUI is integrated into the CardioMat toolbox [26] for cardiac electrophysiology simulations (available at https://github.com/niccolobiasi/CardioMat). A graphically enhanced version is available as a standalone open-source application named AblateTool at 10.5281/zenodo.21335697 to facilitate its distribution in the clinical environment. To assess the efficacy of different catheter ablation strategies in reducing the number of post-ablation residual VTs, our framework enables in silico standardized reinducibility testing after ablation through GPU-accelerated cardiac electrophysiology simulations.

In this work, inspired by the ongoing multicenter clinical trial VOYAGE [27], as a proof-of-concept, we tested the feasibility of our approach for in silico trials by comparing 4 distinct substrate-based ablation approaches in SHD: scar homogenization (SH), primary DZ ablation, primary and secondary DZ ablation, and CMR-guided scar dechanneling (SD). To date, there is still no consensus on the most efficient ablation strategy for SHD patients, and novel approaches are rapidly evolving, including digital twin-guided ablation [28, 16]. In this context, our study has the ambition to provide a tool for in silico comparison of catheter ablation strategies and a first quantitative assessment of emerging ablation strategies, establishing the methodological basis for subsequent large-scale in silico investigations. We considered a cohort of 20 patients, including both ischemic and non-ischemic subjects. For each patient, an anatomically detailed image-based cardiac electrophysiology model was generated from LGE-CMR imaging. Virtual ablation was performed in each VT-inducible patient model by an expert electrophysiologist following each of the 4 considered approaches. Arrhythmia inducibility was reevaluated in each of the 85 computational models to assess the efficacy, burden, and efficiency of each ablation strategy in terms of residual VTs, volume of ablated cardiac tissue, and their ratio, respectively. Our proof-of-concept results highlighted significant differences among the ablation approaches considered, both in terms of efficacy and efficiency. In particular, according to our study, the scar dechanneling approach showed the highest efficiency by significantly reducing the number of inducible VTs while still minimizing ablated volume. On the other hand, scar homogenization is the most effective, but at the cost of very large ablation lesions. This emphasizes the importance of considering the ablated volume when comparing ablation strategies or proposing novel approaches to catheter ablation.

## 2. Methods

### 2.1. Clinical data

10 patients with ischemic cardiomyopathy and 10 with non-ischemic cardiomyopathy undergoing VT ablation with an MRI-aided approach [12] between June 2021 and November 2024 at Pisa University Hospital were retrospectively analyzed. Patients treated up to June 2024 were also enrolled in the prospective randomized VOYAGE trial [27]. The study was conducted in accordance with the principles outlined in the Declaration of Helsinki and was approved by the institutional ethics committee. Written informed consent was obtained from all participants before inclusion in the study. All patients underwent late gadolinium enhancement cardiac magnetic resonance using a 1.5T scanner. In patients with an implantable cardioverter-defibrillator (ICD), wide-band sequences were used to minimize device-related artifacts. LGE-CMR images were processed with the ADAS3D software (ADAS3D Medical, Barcelona, Spain) to analyze left ventricular myocardial enhancement, as previously described [27, 29]. The ADAS processing semi-automatically segments the left ventricle and thresholds the pixel signal intensity (PSI) using customized, patient-specific lower and upper thresholds (35 ± 10% and 52 ± 10% of the maximum signal intensity). The resulting normalized pixel signal intensity (NPSI) ranges from 0 to 1, where NPSI = 0 corresponds to healthy myocardium (PSI below the lower threshold), NPSI = 1 corresponds to core scar (PSI above the upper threshold), and intermediate values (0 < NPSI < 1) represent the border zone. The ADAS output consists of nine NPSI maps, exported as Visualization Toolkit (VTK) files and representing endo-to-epicardial concentric shells from 10% to 90% of the wall thickness. Additionally, ADAS 3D software automatically identifies heterogeneous tissue channels (HTCs) used for implementing the CMR-guided SD virtual ablation approach. HTCs were identified as continuous pathways of border zone tissue, enclosed by scar core or anatomical barriers, that connect two regions of healthy zone tissue [11].

### 2.2. 3D cardiac electrophysiology model generation

The generation of 3D cardiac electrophysiology models incorporating structural and electrical remodeling from ADAS exported VTK shells was described in detail in a previous publication [26]. For the reader’s convenience, we outline the main steps of the process below. For each patient, the ADAS VTK NPSI maps were imported into CardioMat and converted to a single 3D left ventricular geometry associated with a 3D NPSI map (Fig 1). The cardiac geometries were all discretized with a voxel size equal to 250 *um*. All the processing steps were performed in the CardioMat environment, enabling the generation of mesh-free computational models of the cardiac electrophysiology. Ventricular coordinates (i.e., apicobasal, transmural, and rotational coordinates) were computed as defined in [30]. Computation of ventricular coordinates, including rotational, apicobasal, and transmural coordinates, is based on the bi-eikonal normalization method proposed in [31]. A physiologically plausible Purkinje network was generated for each patient according to a constrained optimization method [26, 32]. Myocardial fiber orientation was defined with a rule-based method to account for cardiac anisotropy [33]. Five examples of Purkinje network and fiber orientation assignments for different patients are shown in Fig. 1.

**Figure 1:**
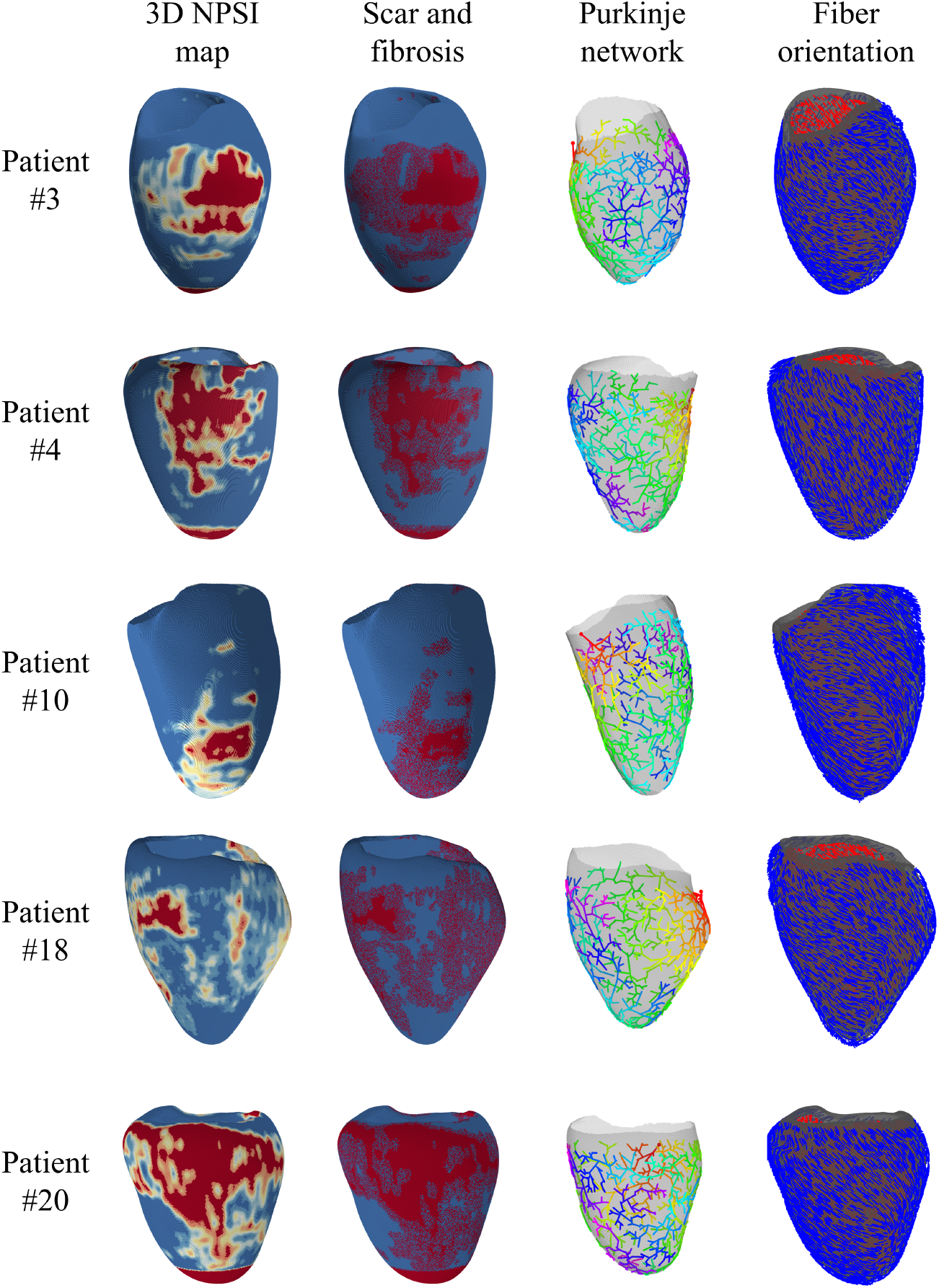
Cardiac electrophysiology models for 5 patients. Each patient is represented on a different row. The first column shows the 3D NPSI maps constructed from ADAS3D shells. The second column shows the assigned scar and fibrotic tissue. Fibrotic and scar voxels are colored in red. The third column shows the Purkinje network, colored according to the CARTO3 colormap (red indicates early activation, purple is late activation). Finally, the fourth column shows fiber orientation. Endocardial fibers are colored in red, whereas epicardial fibers are in blue.

The ionic cellular current was represented with a data-driven phenomenological myocyte model developed by our research group [26, 34]. This choice has two main reasons. First, while maintaining physiologically relevant results, the chosen model showed very high computational efficiency. Given the high number of simulations performed in this study, computational efficiency is a fundamental consideration in the study design. Second, in previous works, we demonstrated that this model is feasible for simulating electrical remodeling in the scar border zone and can generate clinically relevant isochronal maps and DZs [26, 35]. Indeed, myocyte model parameters are dependent on the NPSI, resulting in an increasingly slowed upstroke velocity, reduced action potential notch, and prolonged action potential duration (APD) in the border zone, from healthy cardiac tissue toward the core scar, as reported by experimental studies [36, 37, 38]. This is achieved by linearly relating the myocyte model parameters to the NPSI, as described in [26]. The resulting action potentials for epicardial and endocardial tissues are shown in Fig. 2. Transmural heterogeneity of the cardiac tissue was considered by using the appropriate parameter set in the different layers of the myocardial wall. The endocardial layer was identified as 64% of the myocardial wall, whereas the remaining cardiac tissue was considered epicardium.

**Figure 2:**
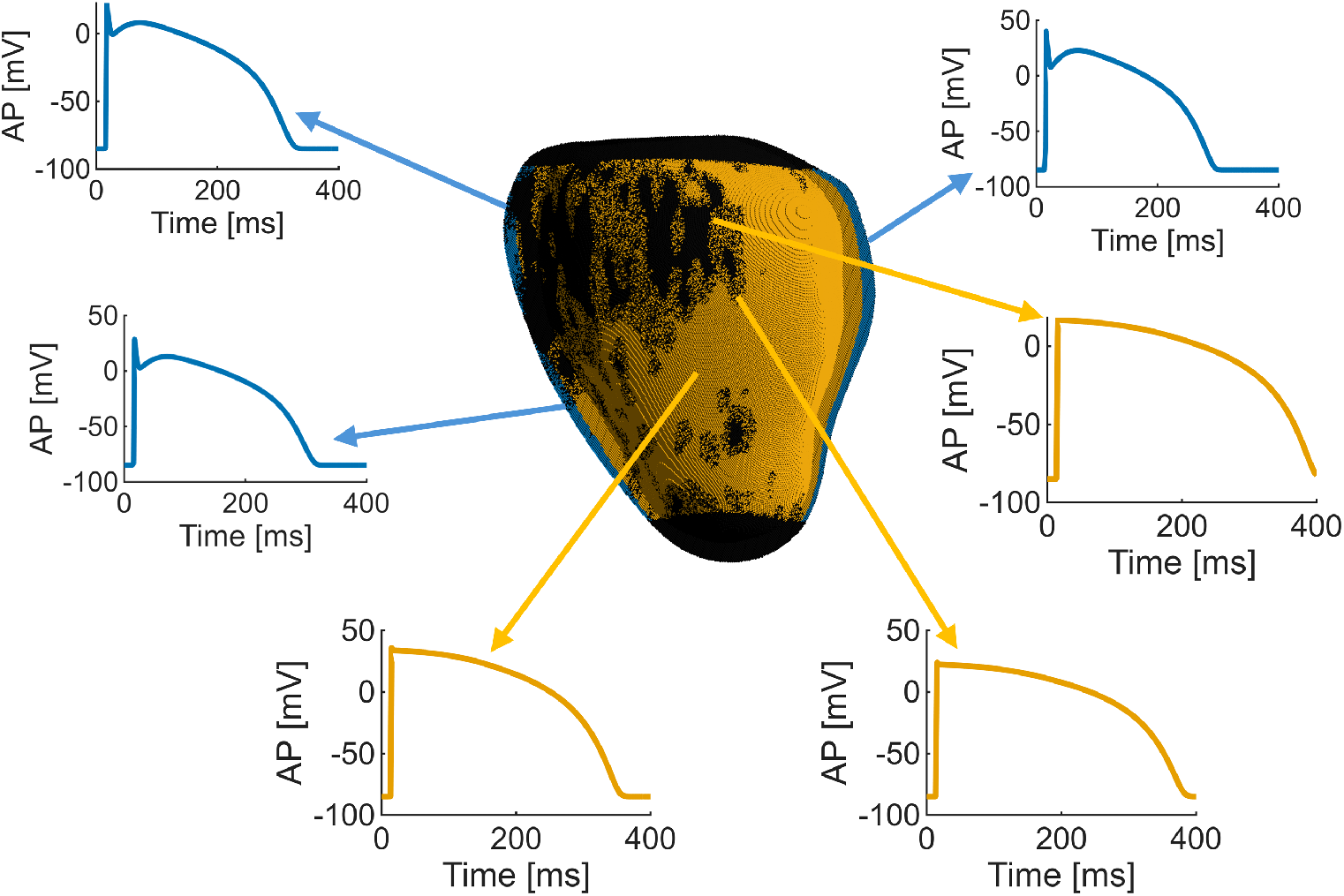
Transmural heterogeneity and infarct-related electrophysiological alterations. The epicardial layer and action potentials are in blue, whereas the endocardial layer and action potentials are in orange. Epicardial and endocardial action potentials were computed on a 2 cm homogeneous cable model with NPSI equal to 0 (healthy), 0.4 (border zone close to healthy tissue), and 0.6 (border zone close to the scar).

We characterized the model behavior and electrophysiological changes for different values of the NPSI based on a 2 cm homogeneous cable model. The maximum APD in the border zone (closest to the scar) was 300 and 376 ms for epicardial and endocardial tissue, respectively. The minimum APD in the border zone was 289 ms for the epicardium and 360 ms for the endocardium. In the healthy epicardial tissue, the APD was 271 ms, whereas in the endocardium, an APD of 334 ms was obtained. Similarly, the maximum upstroke velocity reduces significantly in the proximity of the scar. In the epicardial layer, the maximum upstroke velocity is 209 mV/ms in healthy regions, whereas it varies from 171 mV/ms to 153 mV/ms in the border zone. In the endocardium, the healthy maximum upstroke velocity is 236 mV/ms and reduces to 192 mV/mS in the border zone and up to 171 mV/ms in the proximity of the scar. Note that these values are computed in a 2 cm homogeneous cable model, and may be further altered in a 3D model due to different electrotonic coupling and fibrosis. However, it is worth noting that the electrophysiological properties of the border zone may vary significantly depending on the disease’s stage and may be highly heterogeneous [39, 40], leading to conflicting experimental findings, as for the APD [38]. Additionally, many studies highlighted that structural remodeling is more critical than electrical remodeling in the development of arrhythmias [41, 42, 22].

Based on the 3D NPSI maps, core scar and border zone regions were assigned for each model. Core scar was considered electrically inactive. Patchy fibrosis is commonly observed in the border zones of healed infarcts and is thought to play a significant role in causing local conduction block and triggering arrhythmias [38, 43, 44]. Thus, we accounted for structural remodeling in the border zone by modeling cardiac fibrosis as inexcitable voxels. The density of fibrosis was modulated according to the NPSI, ranging from 40% to 60% in the border zone. These values were selected to align the percentage of fibrotic tissue with the ADAS thresholds. Moreover, a preliminary analysis showed that this choice results in the most arrhythmogenic substrate (see Supplementary material). The fibrosis distribution for 5 patients is shown in Fig. 1.

Simulation of propagating cardiac electrical activity is performed through the monodomain model by using the CardioMat toolbox with a time step equal to 20 *µs*. CardioMat implements an efficient smoothed-boundary GPU monodomain solver based on finite difference discretization on regular grids; thus, no meshing is required. The GPU parallelization implemented in our cardiac electrophysiology solver enables fast cardiac simulations, avoiding the use of supercomputers or clusters. All the simulations reported in this study were executed on two local workstations equipped with NVIDIA GeForce graphics boards (3090 and 5080). Diffusivity parameters were adjusted to achieve conduction velocities of 70 cm/s along the longitudinal axis and 40 cm/s in the transverse direction [45]. In the border zone, the combined effect of fibrosis and electrophysiological remodeling slowed propagation significantly up to about 32 cm/s in the proximity of the core scar. Propagation of action potentials along the Purkinje network was simulated with an Eikonal model with a prescribed conduction velocity of 340 cm/s [46]. The Purkinje network was used only for the simulation of the local activation time (LAT) maps (Section 2.3) and was not considered in the VT induction protocol (Section 2.5). Purkinje-muscular junctions in the border zone and in the core scar were deactivated.

### 2.3. Simulation of activation maps

For each patient model obtained as described in Section 2.2, we carried out a sinus rhythm simulation to compute LATs. Stimulation was delivered in the His bundle, activating the Purkinje network. For each voxel, the activation time was annotated, obtaining a virtual LAT map. LAT was computed as the time instant at which the transmembrane voltage exceeded −20 mV. In fibrotic tissue and core scar, the LAT is not directly available, since transmembrane voltage is not defined; thus, the corresponding LAT value is obtained through nearest neighbor extrapolation. This allows for generating LAT maps with values defined on the whole cardiac surface, similarly to real sinus rhythm electroanatomical mappings. To make the LAT map importable into CARTO 3 systems, we converted the 3D virtual LAT maps into 9 continuous endo-epicardial 2D virtual LAT maps in VTK format through linear interpolation. Similarly, we constructed 8-step isochronal late endo-epicardial maps (ILAM) for easy identification of DZs. Indeed, DZs were defined as regions with 3 isochrones within a 1 cm radius, as described previously [6].

### 2.4. Virtual ablation platform

For in silico catheter ablation, we developed a virtual ablation platform supported by a GUI (Fig. 3) that allows the visualization of the patient’s specific heart using various data visualization modalities. We integrated the virtual ablation platform into a new release of the CardioMat toolbox [26]. To facilitate platform distribution in clinical environments, we also developed a standalone application (i.e., AblateTool) with an enhanced GUI. The AblateTool software is available in open-source from at 10.5281/zenodo.21335697. The only required data for virtual ablation is the CardioMat model associated with a 3D NPSI map, as described in Section 2.2. These can be generated automatically from ADAS VTK shells by CardioMat *adas2vox.m* function and saved to a .mat file. The user can visualize 3D NPSI maps on the patient’s heart anatomy at different transmural levels. The original ADAS VTK files, if available, can be used to visualize ADAS corridors. HTCs identified by the ADAS3D software can be superimposed on the 3D data to facilitate SD ablation. The user can choose whether to visualize the HTCs only relative to the currently visualized transmural layer or all the HTCs. Alternatively, LAT or ILAM data can be visualized at the different transmural levels by reading simulated LAT data generated as described in Section 2.3. The platform allows virtual ablation by simple mouse clicks. At each click, a sphere of tunable size centered at the click point serves as the ablation region. This allows for simulating different ablation modalities, powers, and exposure times. In this work, all the virtual ablations were carried out with an ablation sphere of 1 cm in diameter, resulting in lesions 5 mm deep, as reported for radiofrequency ablation [47]. Each voxel inside the sphere is considered ablated. When the user concludes the ablation procedure, the platform saves a new cardiac model that includes the ablations as scar tissue. An example model is available by default in AblateTool software for testing purposes. The example model is generated starting from a computer tomography image available in the TotalSegmentator dataset [48].

**Figure 3:**
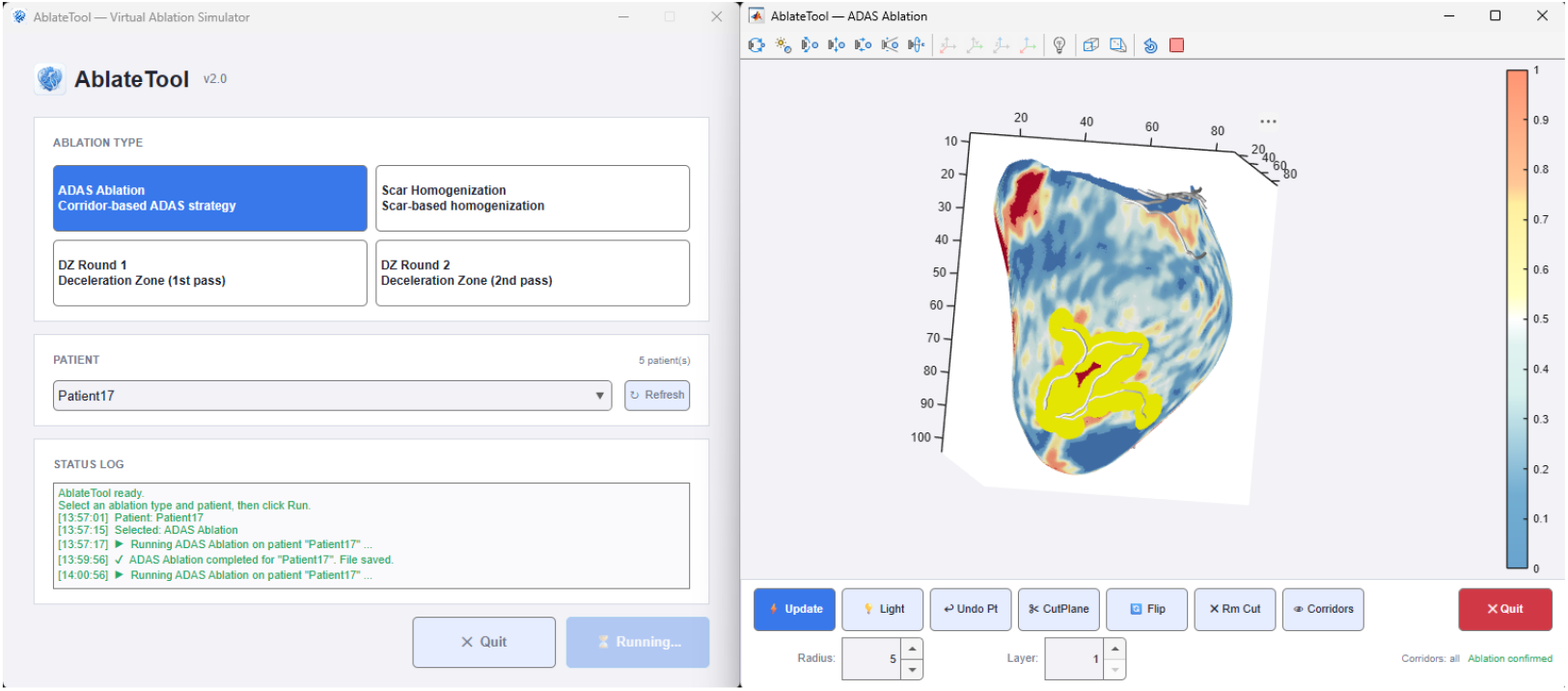
Standalone application for virtual ablation. The figure shows the application during a SD ablation. The ablated regions are colored in yellow. Ablation is performed by simply clicking with the mouse on the 3D model. The camera toolbar at the top of the model and the user interface on the bottom allow for configuring the current view and the size of the ablation lesions.

We employed the virtual ablation platform to perform 4 different ablations in each patient: SH, primary DZ ablation, primary and secondary DZ ablation, and CMR-guided SD. All the virtual ablations were executed endo-epicardially (i.e., by ablating both on the endocardial and epicardial layers when necessary) by an expert cardiac electrophysiologist. Indeed, recent studies suggested that endo-epicardial ablation procedures were associated with a significantly higher success rate and lower VT recurrence [49, 50]. SH ablation was performed on CMR NPSI maps by ablating all the regions corresponding to border zone inside scar areas, first endocardially and then epicardially. During the procedure, particular care was given to avoid the creation of iatrogenic corridors between ablation lesions. In regions with a thin myocardial wall, an ablation lesion positioned endocardially may also have an effect epicardially and vice versa. Thus, after epicardial ablation, the endocardial layer was checked again for the generation of corridors between ablation lesions. CMR-guided SD ablation was performed by visualizing the HTCs as identified by ADAS3D and ablating at the ends or along each HTC in order to interrupt conduction along the channel. Eventual new iatrogenic corridors between ablation lesions were also ablated to avoid procedure-related arrhythmias. Primary DZ ablation targeted the DZs identified by ILAM as regions with at least 3 isochrones within a 1 cm radius [6]. For patients with residual VT inducibility, secondary DZs were identified by generating new LAT data from the cardiac model resulting from the primary DZs ablation procedure. A new ablation procedure was then performed on this model, targeting secondary DZs.

### 2.5. Virtual VT induction protocol

For each patient, we performed 17 simulations. In each simulation, we applied a standardized stimulation protocol in the centroid of one of the 17 left ventricle AHA segments (Fig. 4). The AHA segments of the LV were identified by thresholding the apicobasal and rotational coordinates, which are automatically generated during the model construction. Stimulation was modeled by applying a transmembrane current of 50 *mV/ms* within a 1 *cm* diameter sphere at each AHA segment individually. A sequence of five stimuli with a basic cycle length of 600 ms was followed by three extrastimuli applied at the same anatomical site (S1 600 ms x 5, S2 350 ms, S3 300 ms, S4 280 ms). An automated procedure was used for the implementation of the inducibility test. First, in the prepacing stage, the 5 S1 stimuli were delivered. If the stimulation region is completely superimposed on scar tissue, the simulation was skipped, and the program moves to the next AHA segment. At the end of the prepacing stage, the final state of the system was saved to a .mat file and used as an initial condition for the inducibility stage. This allows for saving computational effort when different S2 inducibility protocols are tested. However, unlike the original VARP protocol [21], we used a fixed number of extrastimuli to make the comparison between ablation strategies in terms of residual VTs more robust, ensuring that the same stimulation is applied before and after ablation. Such an aggressive stimulation protocol was selected to highlight potential differences between ablation strategies. VT was considered induced if electrical activity lasted at least 1 second after the end of the stimulation protocol. In such cases, a checkpoint file was saved, and VT was then simulated for additional 4 seconds. If electrical activity was still present 5 seconds after the end of the stimulation protocol, the simulation was labeled as sustained VT. The virtual VT induction protocol was then repeated on the ablated models. For each patient and each ablation strategy, we recorded the total number of induced VT episodes and sustained VT episodes. This process involved 85 simulations per patient, each potentially lasting up to 8.33 seconds (3.3 seconds of stimulation protocol and 5 seconds for arrhythmia monitoring). Simulations in which VT was neither inducible nor sustained were terminated earlier, once all voxels had returned to the resting state. Notably, all the induction testing is completely automated. An example script replicating the VT induction protocol is available at the CardioMat Zenodo repository (https://zenodo.org/records/17398365).

**Figure 4:**
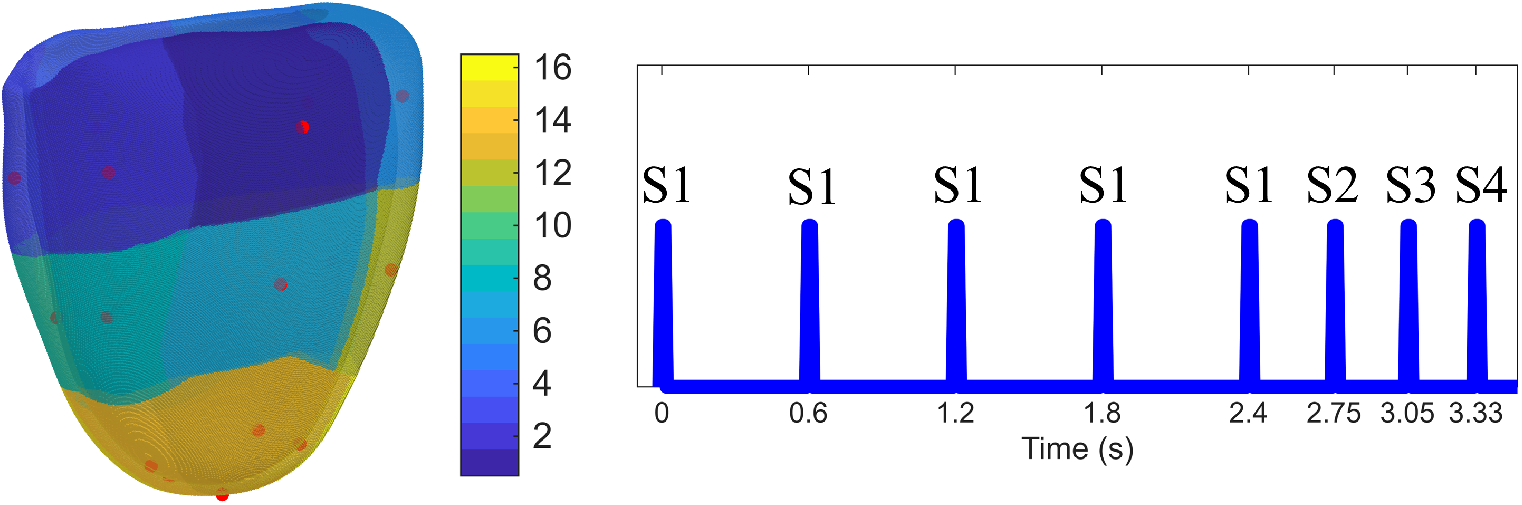
Stimulation protocol adopted for the induction of VT in the virtual patients. Stimulation was delivered at the centroids (red dots) of AHA segments. On the left, AHA segments are colored according to the colorbar. The right side of the figure shows the stimulation protocol (S1 600 ms x 5, S2 350 ms, S3 300 ms, S4 280 ms), which was applied separately at each AHA segment.

The last two seconds of sustained VT simulations are saved to a binary file for subsequent visualization and analysis with a time resolution of 5 ms. For each sustained VT, the cycle length was computed through a pseudo-ECG computation by assuming an infinite and homogeneous volume conductor approximation, and a simulated LAT map was generated to better visualize the VT circuit. LAT was computed by tracking voxel activation in the time window between the last two R peaks of the pseudo-ECG. As in the sinus rhythm case, LAT was linearly interpolated in the fibrotic voxels and converted to CARTO-importable continuous endo-epicardial 2D virtual LAT maps in VTK format. To enhance VT circuit visualization in these LAT maps, the scar core was shown in white. A specific function (*lat2shell.m*) is made available in CardioMat for creating CARTO-importable LAT maps from simulated LAT data. All the induced sustained VT circuits were subsequently analyzed and visually inspected by an expert electrophysiologist. VTs sharing the same circuits were identified, and the number of unique VTs and total VTs were counted for each patient. Unstable VTs and ventricular fibrillation patterns were always classified as unique. To assess and compare the efficacy and efficiency of the different ablation strategies, we mainly focus on unique VTs, which quantifies the number of possible reentry circuits that could establish on a patient’s heart, independently of the number of times they are observed. The use of simulated unique VT circuits as an indicator of a patient’s VT inducibility has already been supported by previous studies assessing VT risk with cardiac electrophysiology models [51, 52]. Differences among ablation strategies were assessed using non-parametric tests followed by post-hoc pairwise comparisons on relevant metrics quantifying ablation efficacy, burden, and efficiency. Efficacy was quantified by the number of unique sustained VTs and the percentage of ablated VTs relative to baseline (i.e., the number of VTs before ablation). Ablation burden was measured as the absolute and relative ablated volumes. Efficiency was quantified by the number of ablated VTs per milliliter of ablated volume and by the percentage of ablated VTs normalized by the relative ablated volume (i.e., the percentage of suppressed VTs achieved by ablating 1% of the myocardial tissue). None of the analyzed variables followed a normal distribution; thus, they are presented as medians [25*^◦^ −* 75*^◦^* interquartile ranges, unless otherwise stated]. Since simulated data were paired across ablation strategies on the same patient, differences among conditions were assessed using the Friedman test. When a significant overall effect was detected, post-hoc pairwise comparisons were performed using Wilcoxon signed-rank tests with Bonferroni correction applied to adjust for multiple comparisons.

## 3. Results

### 3.1. VT inducibility

Inducibility simulations were correctly executed in the 20 patient-specific models. Sustained VT was inducible in 17 out of 20 patients, while two ischemic patients and one non-ischemic patient were non-inducible. Overall, 127 sustained VTs were induced, 63 in the 8 ischemic patients and 64 in the 9 non-ischemic subjects. However, only 88 of the total number of VTs corresponded to unique reentrant circuits (46 in ischemic patients). This resulted in an average of 4.4 unique VTs per patient (5.2 per inducible patient), consistent with previous computational studies [51]. There is no significant difference in terms of VT inducibility between ischemic and non-ischemic patients. The average cycle length of induced VT was 390 ms (std *±*60 *ms*), with a minimum of 310 ms and a maximum of 657 ms, comparable with the clinical range [53]. Average cycle length did not differ between ischemic and non-ischemic patients (384 *±* 53 *ms* vs 393 *±* 65*ms*).

Figure 5 shows a simulation in which sustained arrhythmia is successfully induced. After the last extrastimulus, propagation fails in highly fibrotic regions close to the core scar due to the impaired conduction. Thus, these regions remain excitable. The fibrosis induces the action potential wavefront to curl, newly exciting the healthy tissue and establishing reentrant activity. After some unstable reentries, the action potential wavefront starts to rotate around a scar region steadily. To better visualize VT circuits, we generated a simulated electroanatomical mapping of the VTs. We observed four main types of VT circuits, depending on the anatomical structures involved and the geometrical features of the reentry circuit. A representative example of each VT circuit type is shown in Fig. 6. The core scar was colored in white to delineate the VT circuit. In the first example, the VT circuit is established around the core scar (Fig. 6A), whereas in the second one, the circuit involves HTCs on a single transmural shell (Fig. 6B). These also include the so-called incomplete reentries, which cannot be visualized entirely from endocardial and epicardial surfaces due to the presence of intramural circuits [54]. Interestingly, arrhythmogenic HTCs may also be located between scar tissue and the mitral valve openings. On the other hand, the third type of VT circuit is confined completely to the border zone and is induced by the fibrotic tissue (Fig. 6C). In some cases, none of the previous VT patterns can be visualized on a single transmural shell (even in the midmyocardium), highlighting the presence of a 3D circuit with the isthmus located transmurally (Fig. 6D). Thus, the types of VT we observed are coherent with the hyperboloid structure of the VT circuit recently proposed by Nishimura et al. [55]. Additionally, ventricular fibrillation patterns can be observed in patients with large border zones (Fig 7). In this case, a clear reentry circuit and an associated cycle length cannot be identified.

**Figure 5:**
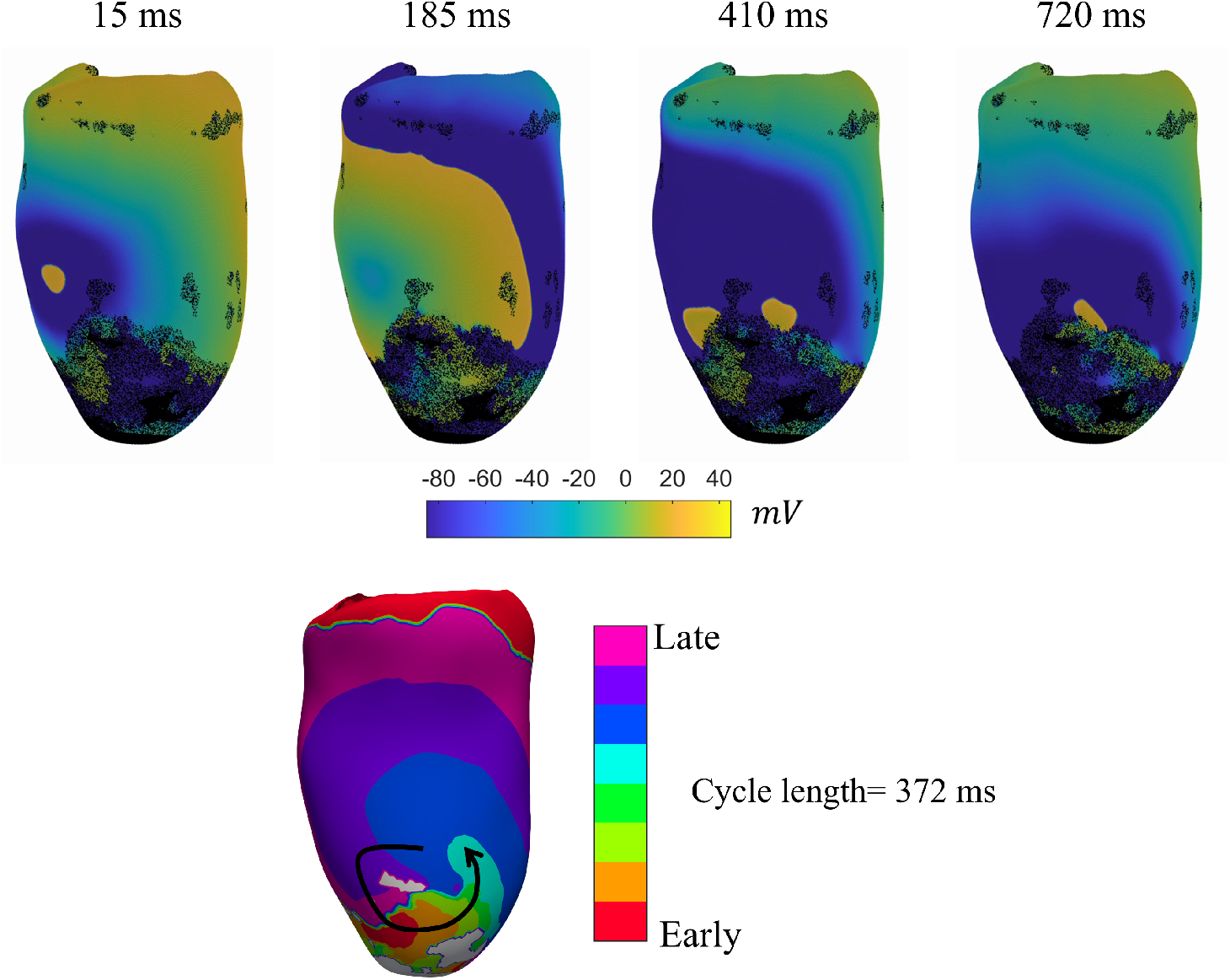
Induction of VT in patient 6 with decremental pacing applied at segment 12. Four snapshots of transmembrane potential maps are shown, each corresponding to a different time instant. Time instants are referred to the last stimulation of the pacing protocol. On the bottom, the final stabilized circuit is shown with the associated cycle length.

**Figure 6:**
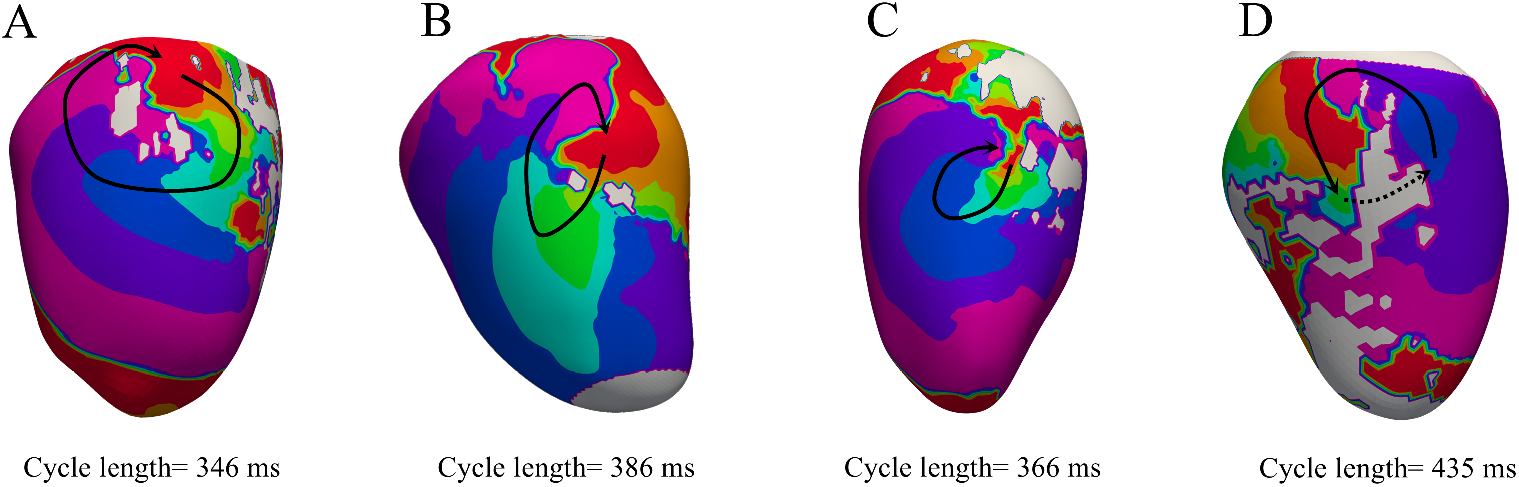
Different types of observed reentry circuits. A) Reentry around the core scar (subject 16). B) Reentry through an HTC in the epicardium (subject 20); C) Border zone reentry induced by fibrosis (subject 5); D) 3D reentry circuit with isthmus located transmurally (subject 18). CARTO3 colormap was used. Black arrow highlights the reentry circuit

**Figure 7:**
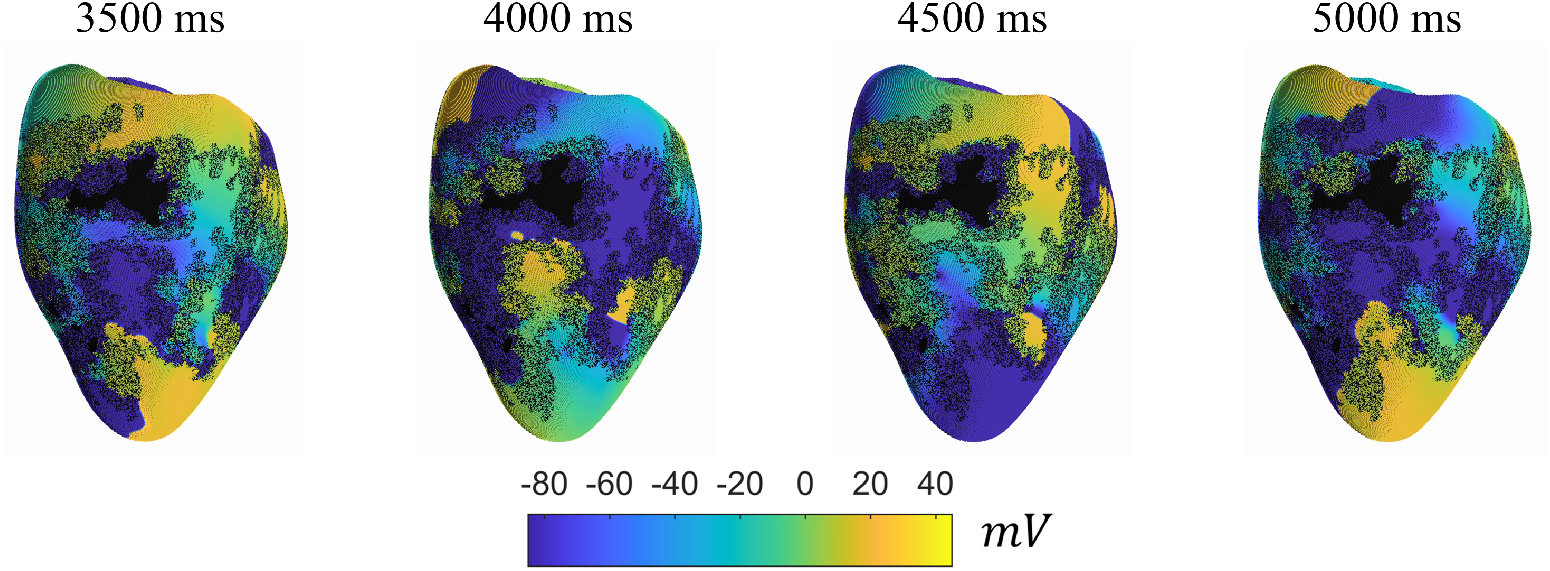
Simulation showing a ventricular fibrillation pattern on patient 17. No clear and sustained reentry circuits can be identified. Time instants are referred to the last stimulation of the pacing protocol.

### 3.2. Simulated ILAM maps and virtual ablations

Sinus rhythm LAT and ILAM maps were correctly simulated, and virtual ablations were successfully conducted in all the 17 VT-inducible patients by a clinical electrophysiologist using the standalone AblateTool software. Fig 8 shows some examples of ADAS-derived NPSI maps, simulated ILAM maps, and virtual ablations for different patients and ablation strategies. Late activation and DZ regions in ILAM maps are mostly superimposed with core scar and border zone tissue identified by CMR analysis, in alignment with clinical observation [56]. CMR-guided SD ablation focuses on the corridors identified by ADAS3D. Depending on the patient’s structural substrate, the ablation volume can be quite large related to the total volume of myocardial tissue. SH ablation covers the whole border zone, homogenizing it with core scar. In this way, the border zone impaired tissue will no longer be able to conduct electrical impulses and thus sustain arrhythmias. Primary DZs identified in ILAM maps usually cover a small amount of the cardiac tissue in the regions close to the core scar tissue. Consequently, primary DZ ablation is in most of the cases very conservative. After primary DZ ablation, remapping always showed additional DZs (i.e., secondary DZs) in locations adjacent but distinguished from the original primary DZs. As a result, secondary DZ ablation enlarges the original ablation.

**Figure 8:**
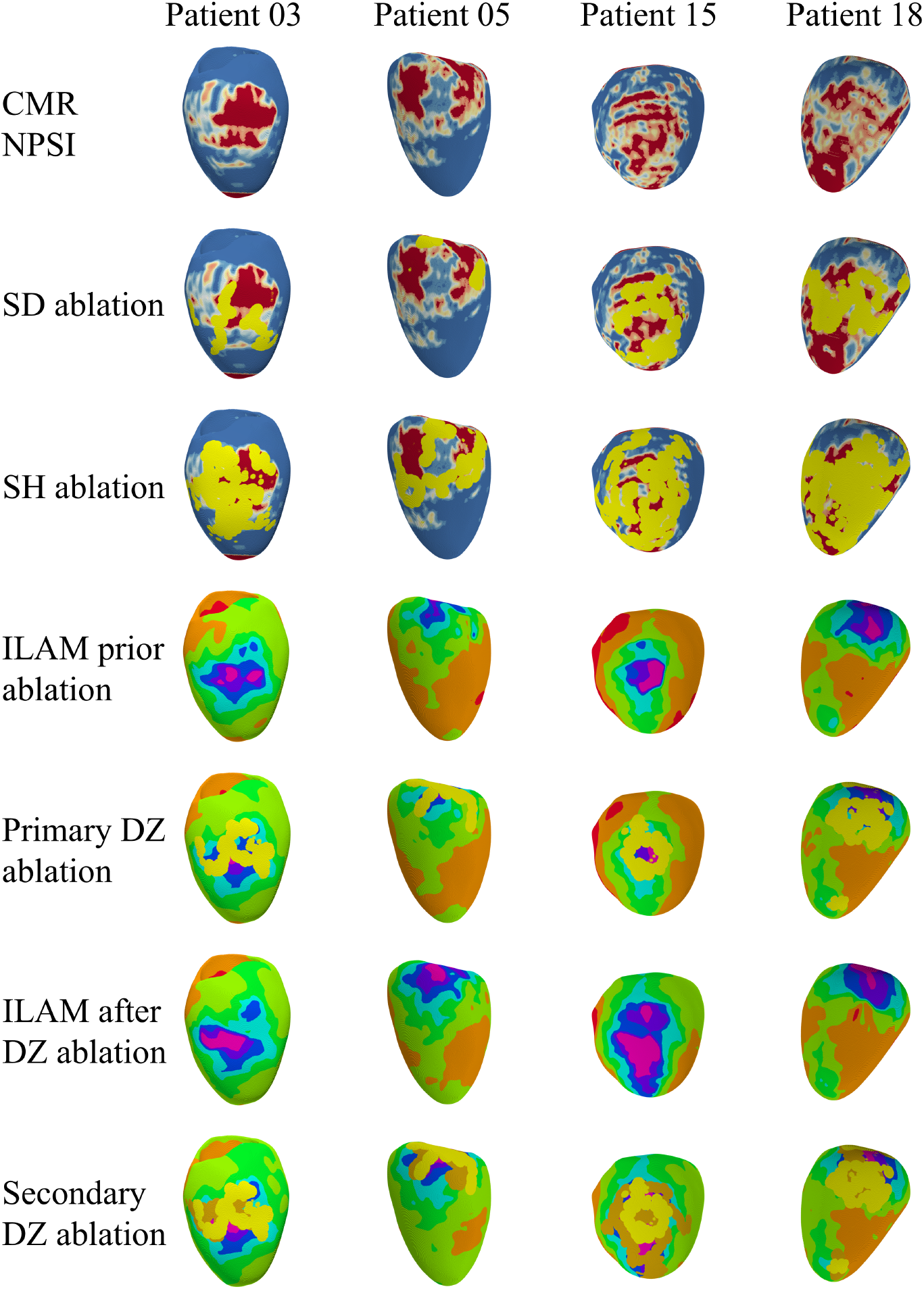
Examples on four patients (3, 5, 15, and 18) of simulated isochronal mappings and virtual ablations performed with the AblateTool software. Standard ADAS colormap was used for CMR NPSI visualization. Isochronal mappings (ILAMs) are colored according to the CARTO3 colormap. Ablated regions are colored in yellow; only secondary DZs ablation lesions are colored in dark yellow.

Substantial differences were observed and statistically highlighted in terms of ablated myocardial volume (Fig. 9). Overall, the SH approach is the most aggressive ablation strategy, covering, on average, 21% of the heart volume (about 20 ml), which is significantly higher than all other approaches (median: 19.61 [12.02-27.91] ml, *P <* 0.01). On the other hand, the most conservative ablation strategy among the tested ones targets the primary DZs, resulting, on average, in ablation of about 8% of myocardial tissue (7 ml), which is significantly lower than all other approaches except the CMR-guided SD approach (median: 7.1 [4.91-8.46] ml, *P <* 0.01). The further ablation of secondary DZs significantly increases the ablated volume to 11 ml on average, covering about 12% of the myocardial tissue (median: 9.45 [7.26-15.35] ml, *P <* 0.001). Similarly, CMR-guided SD ablation covers, on average, 10% of the myocardial tissue, corresponding roughly to 9 ml (median: 8.96 [6.18-13.47] ml). No further statistically significant differences were observed regarding the ablated volume. When considering the ablated volume normalized to the total myocardial volume, the same statistically significant differences were detected.

**Figure 9:**
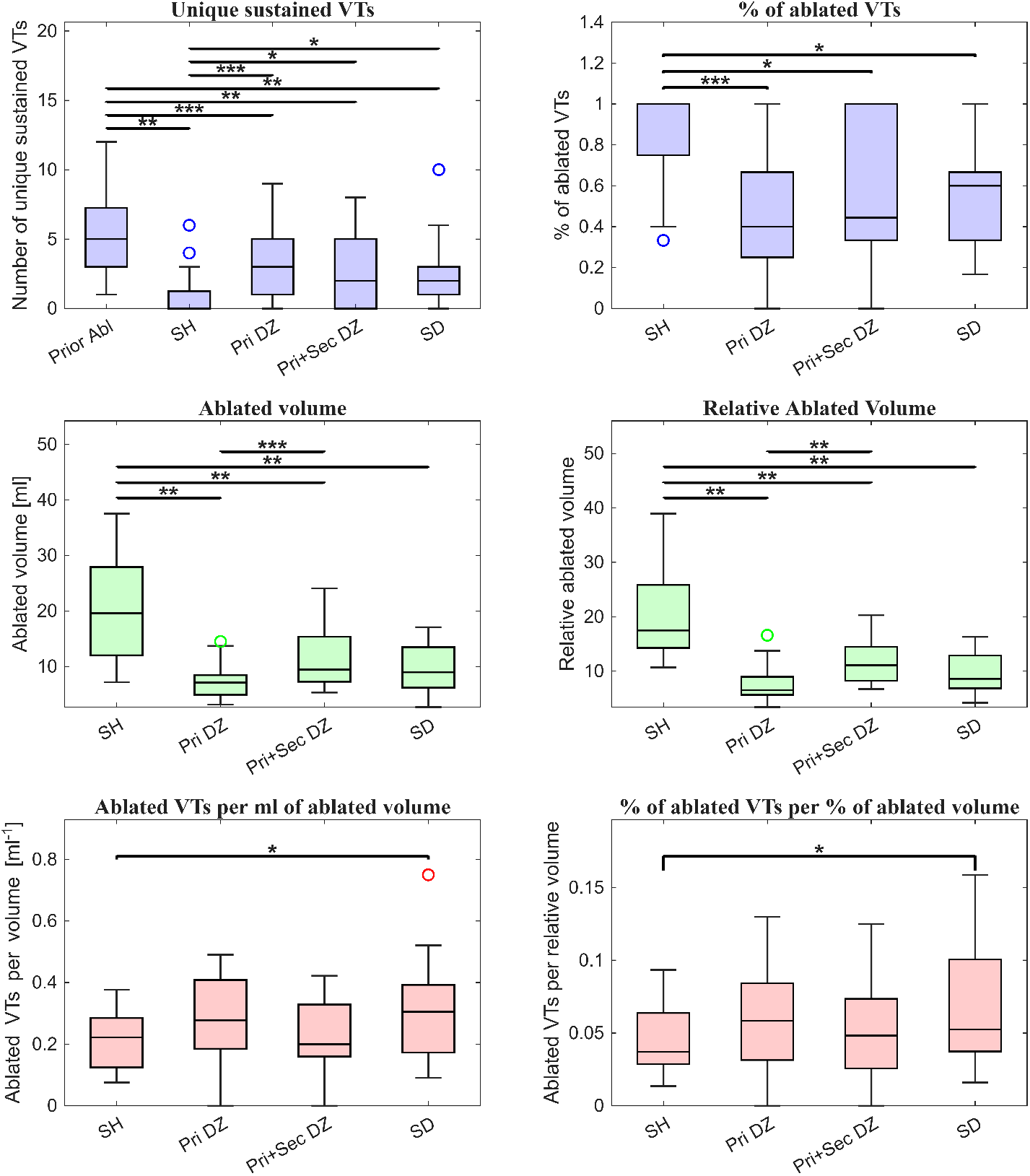
Efficacy, ablation burden, and efficiency of the tested virtual ablation strategies. Boxplots report the number of unique sustained VTs, the percentage of VTs suppressed relative to baseline, the absolute and relative ablated volume, the number of suppressed VTs per milliliter of ablated tissue, and the percentage of suppressed VTs normalized by the percentage of ablated myocardial volume. The compared conditions include baseline before ablation (Prior Abl), scar homogenization (SH), primary deceleration-zone ablation (Pri DZ), primary plus secondary deceleration-zone ablation (Pri+Sec DZ), and CMR-guided scar dechanneling based on ADAS3D corridors (SD). Boxes represent the interquartile range, horizontal lines indicate medians, whiskers show non-outlier ranges, and circles denote outliers. Statistical comparisons were performed using Friedman tests followed by Bonferroni-corrected Wilcoxon signed-rank tests. \**P <* 0.05, \*\**P <* 0.01, \*\*\**P <* 0.001.

### 3.3. Post-ablation VT induction

Efficacy and efficiency of the four considered ablation strategies were compared in terms of unique sustained VT, percentage of ablated VT relative to the baseline, absolute and relative ablation volume, ablated VT per ml of ablation volume, and percentage of ablated VTs per percentage of ablated volume. The analyzed data derived from our in silico trial are reported in Table 1 of the Supplementary Material. All four ablation strategies significantly reduced VT inducibility compared with baseline (Fig. 9). SH achieved the largest reduction, with an average of 1.05 residual VTs per patient (*P <* 0.01). More than half of the patients were not inducible after SH ablation (residual VT median: 0 [0-1.25]) CMR-guided SD reduced significantly inducible VTs to 2.52 per patient (median: 2 [1-3], *P <* 0.01). Primary DZ ablation resulted in 3.5 residual VTs per patient (median: 3 [1-5], *P <* 0.001), while the addition of secondary DZ ablation further reduced inducible VTs to 2.92 per patient (median: 2 [0-5], *P <* 0.01). SH ablation was significantly more effective than all the other ablation strategies (*P <* 0.001 vs primary DZs ablation, *P <* 0.05 when compared to the other strategies). No other statistically significant differences in VT suppression were observed among the four strategies. When considering the percentage of ablated VT, the same significant differences are observed. SH ablation reduces VT events by 85% on average (median: 100 [75-100] %), whereas primary DZs ablation suppresses 44% of VTs (median: 40 [25-66.67] %). Targeting secondary DZs improves VT ablation to 59% (median: 44.44 [33.33 100]). CMR-guided SD ablation suppressed 57% of VTs (median: 60 [33.33 66.67] %)

When VT suppression was normalized by ablated tissue volume, CMR-guided SD showed the highest efficiency, significantly better than SH(*P <* 0.05), and achieved substantial reductions in VT inducibility while minimizing myocardial loss. Indeed, CMR-guided SD suppresses, on average, 0.31 VTs per ml of ablated myocardial tissue (median: 0.31 [0.17-0.39] VTs/ml). In contrast, whereas SH remained the most effective strategy in absolute terms, its efficiency was the lowest (median: 0.22 [0.12-0.28] VTs/ml) among the tested ablation strategies, due to the extensive ablation volume. Both primary (median: 0.27 [0.18-0.41] VTs/ml) and primary+secondary DZs (median: 0.20 [0.16-0.33] VTs/ml) ablation did not differ significantly from SH and CMR-guided SD ablation strategies. The inclusion of secondary DZs as ablation targets shows a trend toward reduced efficiency, moving from an average of 0.28 VTs/ml to 0.23 VTs/ml. When considering the normalized version of the efficiency indicator, we still observed significantly higher efficiency of CMR-guided SD approach when compared to SH ablation.

### 3.4. Mechanism of VT ablation: success or failure

Figure 10 shows the results of every induction test performed on patient 10, considered as a case study to assess the mechanisms leading to success or failure of VT ablation. The patient’s model was inducible from 8 AHA segments before ablation. After SH ablation the model was completely non-inducible, with only one non-sustained reentry observed. Primary DZs ablation suppressed the majority of VT reentry circuits; however, sustained reentry can still be induced by two distinct AHA segments. Considering, for example, the VT induced from segment 11, the isochronal mapping reveals that while the original circuit was indeed suppressed, catheter ablation introduces a new epicardial corridor that serves as a reentry circuit. Indeed, primary DZs ablation was performed looking at ILAM maps, without considering scar imaging. Further ablation of secondary DZs definitely suppressed all sustained VTs. CMR-guided SD results in a model inducible from segments 2 and 11. Whereas pacing from segment 11 induced a VT also in the baseline model, sustained VT induced from segment 2 was not observed in the baseline model. The endocardial ablation of ADAS-derived corridors generates a completely new reentry circuit in the midcardium that was not observable either from the epicardium or the endocardium. Figure 11 explains how catheter ablation impedes generation of reentrant activity by showing the induction protocol prior to and after CMR-guided SD ablation. Basically, catheter ablation replaces arrhythmogenic fibrotic border zones with completely inexcitable dense scar, which is no longer able to conduct electrical impulses. Additionally, even if in several cases a non-sustained reentry can be observed even after ablation, the dense scar impedes the establishment of stable reentry circuits by splitting closed-loop pathways. Thus, electrical activity annihilates after a few rotations.

**Figure 10:**
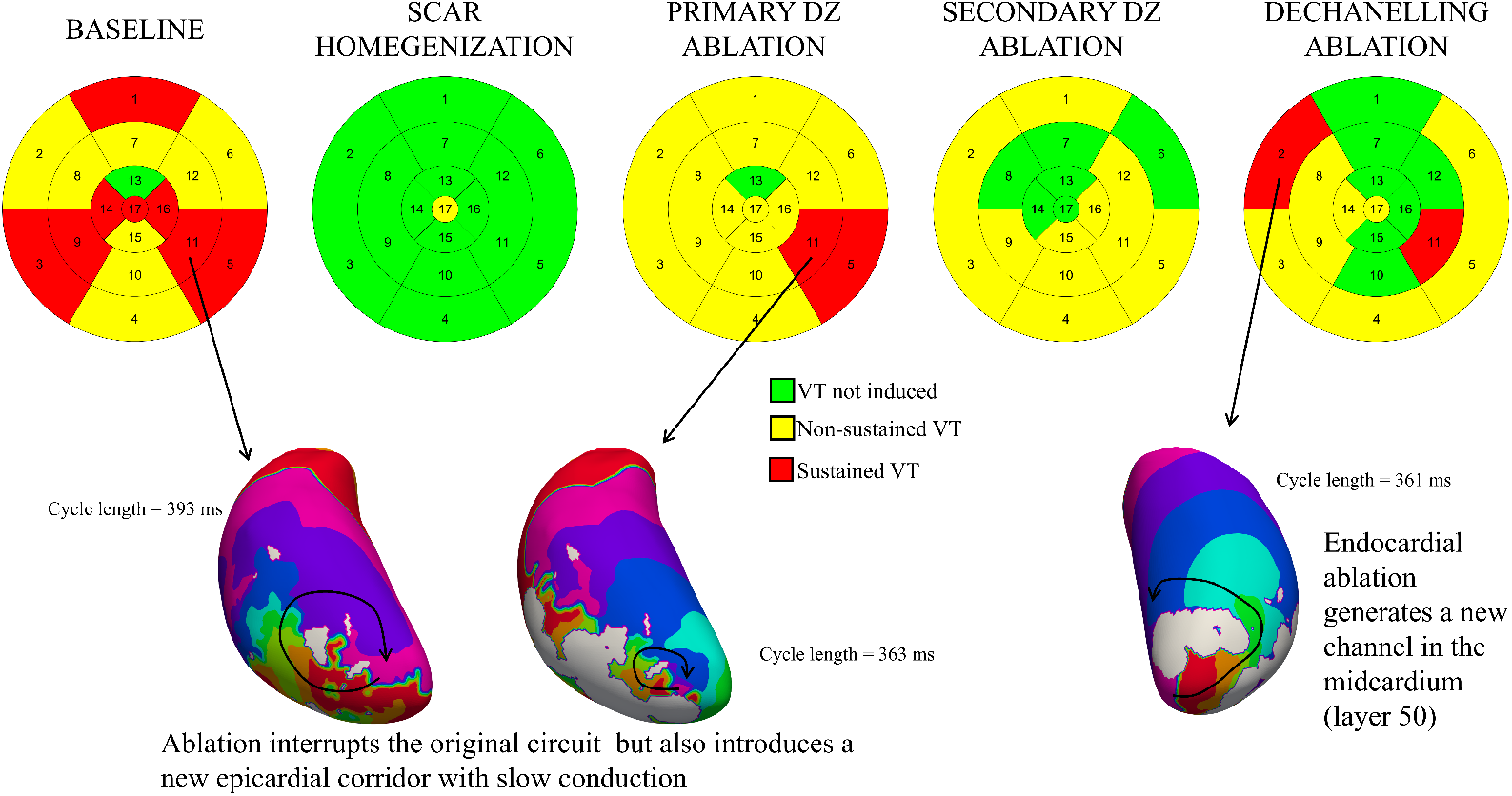
Patient 10 case study assessing VT inducibility before and after ablation. Bullseye plots show AHA segments from which VT is induced. Red segments indicate that sustained VT was induced from those segments, whereas yellow segments represent non-sustained VT. Green segments indicate that no VT was induced. Three representative reentry circuits are shown to highlight different mechanisms of ablation failure. The cycle length of each VT was reported on the side of the VT LAT map.

**Figure 11:**
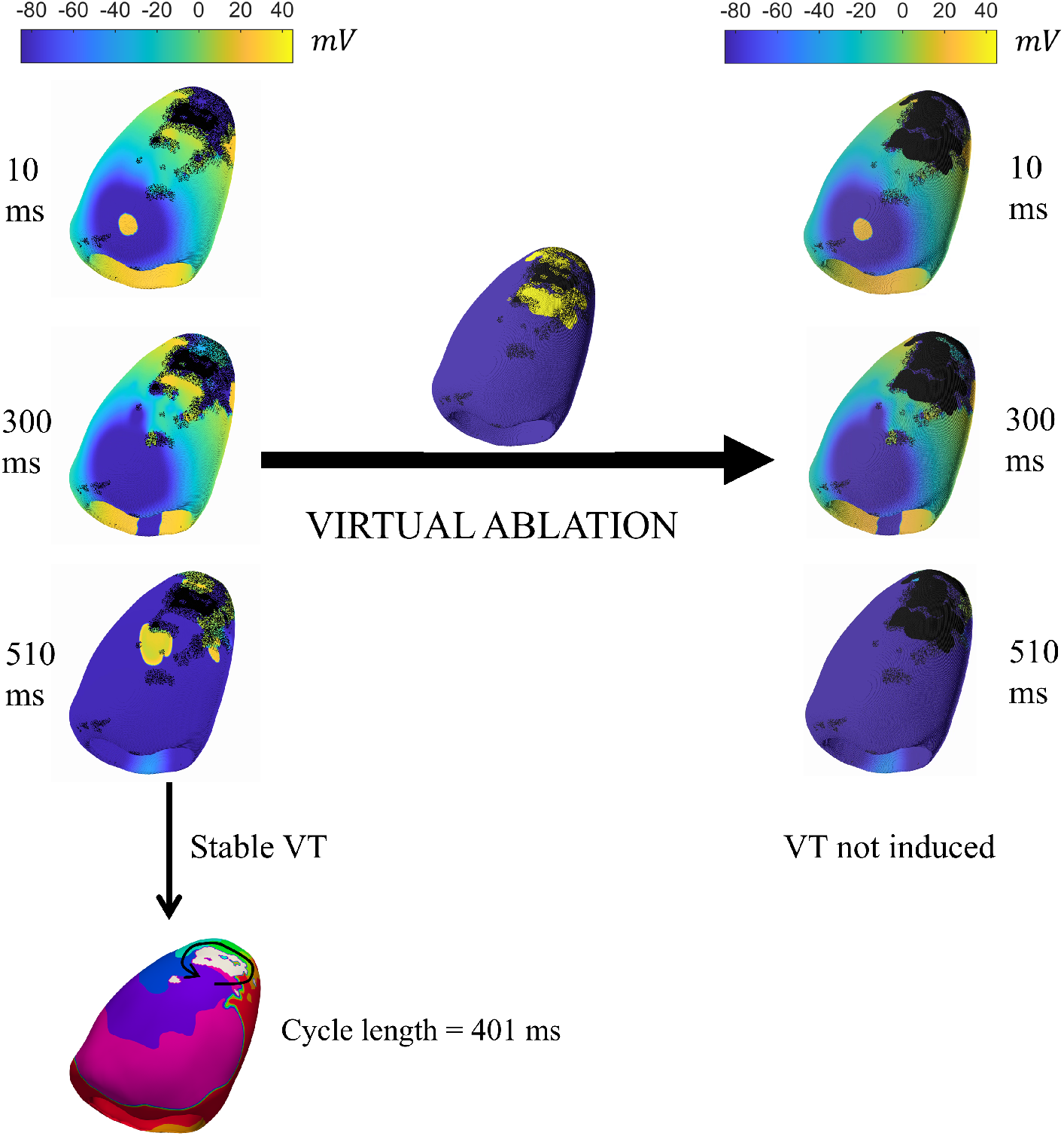
Example of successful VT ablation with SH strategy on patient 10. VT was induced from segment 1 prior to ablation. After the virtual ablation shown in the center of the figure (yellow regions), VT was not inducible.

## 4. Discussion

The primary contribution of this work is an image-based computational framework for conducting controlled in silico trials of substrate-based VT ablation strategies. Starting from clinical imaging data, the framework generates patient-specific electrophysiological substrates, supports interactive virtual ablation, and enables standardized reinducibility testing across different ablation strategies. As a proof of concept, we applied the framework to compare four clinically relevant strategies inspired by current substrate-based VT ablation approaches. However, our proposed framework could be employed for assessing the efficacy and efficiency of any catheter ablation approach, including novel and not yet clinically used innovative strategies, with respect to both the baseline model and standard ablation procedures. Indeed, any virtual lesion set can be introduced into the image-derived substrate and evaluated using the same standardized reinducibility protocol.

The results obtained by the proof-of-concept in-silico clinical trial enabled by the framework presented in this paper suggested that SH ablation is more effective than limited, localized ablations. This finding is in line with the VISTA clinical trial [5], which concluded that extensive substrate ablation is superior to ablation targeting only clinically observable and stable VTs. Nevertheless, our results also suggest that SH ablation targets an extensive volume of myocardial tissue, significantly higher than all other strategies. This finding highlights the importance of assessing VT ablation approaches (both clinically and in silico) not only in terms of efficacy (e.g., VT-inducibility, VT-free time interval post-ablation) but also in terms of efficiency, giving an important weight to the ablated volume. Indeed, considering the extreme case, an ablation covering the whole heart volume will result in a completely VT-free model, but this ablation will clearly not be acceptable. Assessment and quantification of the efficiency of a catheter ablation approach is particularly important when novel strategies are proposed, including digital-twin-guided ablation [28, 18]. Thus, we introduced a novel indicator for quantifying the efficiency of a virtual catheter ablation procedure by normalizing the number of ablated VT circuits to the ablated volume. Considering our newly defined efficiency indicator in both absolute and normalized versions, CMR-guided SD was shown to be significantly more efficient than SH ablation. The conducted in silico trial also suggested that DZs ablation and CMR-guided SD achieve comparable results, in line with the study of Vazquez-Calvo et al. [57], which has previously demonstrated that DZs show high correlation with CMR-derived corridors. Thus, the similar performance of a DZ-based and an image-based ablation strategy indirectly highlights the reliability of our modeling approach in predicting DZs. The inclusion of secondary DZs did not bring a significant improvement in terms of efficacy, while the amount of ablated volume is significantly higher. Nevertheless, these results may be affected by the limited sample size, preventing to observe significant differences between the two strategies in terms of efficacy or efficiency. Considering the percentage of ablated VT, targeting secondary DZs seems to improve the efficacy of ablation, but this should be confirmed on a larger sample size.

Importantly, our framework for in silico comparison of catheter ablation strategies is independent of the modeling assumption made in the conception of this study. For example, in our proof-of-concept study, fibrosis density in the border zone was modulated according to the NPSI and ranged from 40% to 60%. This choice was intended to make the amount of inexcitable tissue consistent with the ADAS thresholding and to reproduce conduction slowing and discontinuous propagation in the heterogeneous border zone. The proposed framework is not restricted to this specific fibrosis representation and can accommodate alternative structural remodeling models. Different assumptions can be made regarding border zone remodeling. Indeed, our approach in modeling SHD substrate differs significantly from previous works (e.g., [15, 17, 20]). Conduction slowing in the border zone is usually obtained by modifying the myocardial conductivities in the border zone. Differently, we focus our computational models on cardiac fibrosis, which causes conduction slowing and wavefront fragmentation in alignment with electrogram QRS fragmentation reported in the border zone of both ischemic and non-ischemic patients. The observed mechanism of VT initiation is strongly linked to the presence of fibrosis and is aligned with percolation theory [58]. Indeed, previous works have hypothesized the role of relatively small structural abnormalities in VT initiation and maintenance [59, 60]. However, the majority of previous works focusing on VT secondary to SHD overlooked the role of cardiac fibrosis. Consequently, the VT initiation was mainly described as the consequence of a larger conduction block and the immediate establishment of a stable reentry circuit anchored to scar tissue. Additionally, the use of three extrastimuli, while improving identification of VT circuits, may hide other VT initiation mechanisms, more similar to those observed in [15] Similarly, our framework could in principle support different imaging modalities capable of characterizing the arrhythmogenic substrate. In this proof-of-concept study, LGE-CMR-derived ADAS3D maps were used to generate the patient-specific substrates and to carry out CMR- guided ablation approaches; however, patient-specific structural substrate could also be obtained from CT imaging by processing wall thickness [51]. The inclusion of CT data in the pipeline would also allow the testing of CT-guided ablation approaches [10].

Additionally, the AblateTool software can be used independently from the CardioMat solver and coupled with different electrophysiological simulators. For example, the *vox2carp.m* function contained in the CardioMat repository allows exporting the anatomical model, including the structural substrate and the ablation, into the openCARP format [61]. This feature also enables the use of different open-source frameworks for VT-inducibility testing, such as AutoVARP [62].

Our framework for in silico trial of catheter ablation strategies presents some important limitations. First, as in recent works [17, 28, 22, 20], we used the same baseline electrophysiological parameters for each patient, without including a personalization process based on electrophysiological data, such as those presented in [63, 64]. All the models were constructed and personalized only on structural data (i.e., LGE-CMR). However, the focus of this work was the development of a framework for in silico trials of catheter ablation strategies rather than predicting patient-specific risk or optimal therapy, making this limitation less relevant to our study. The Purkinje system was included for sinus rhythm activation and virtual electroanatomical mapping but not for VT induction. Therefore, the present study specifically focuses on myocardial scar-related reentrant mechanisms and does not address Purkinje- mediated ventricular arrhythmias. However, it is worth mentioning that the Purkinje system may play a significant role in the induction and maintenance of VT also in SHD patients [65, 66].

Regarding the proof-of-concept study presented, the main limitation is represented by the small sample size, which did not allow a comprehensive characterization of different ablation strategies and particularly their separate assessment in ischemic vs non-ischemic patients. Furthermore, due to computational constraints, VT was observed only for 5 seconds, whereas clinical observation usually lasts around 30 seconds. Thus, our study could have overestimated the number of sustained VTs, since with a longer observation interval, some VTs classified as sustained could annihilate after 5 seconds. Additionally, to maintain the study computationally tractable, we employed a phenomenological model for representing ionic current at the myocyte membrane. This choice was motivated by the large number of simulations performed. With the actual configuration, 1 s of simulation requires, on average, about 450 seconds to be completed on a NVIDIA GeForce 3090 GPU (and slightly more for the 5080 GPU), varying with the size of the patient’s heart. Our study involved 1445 simulations, each potentially lasting up to 8.33 seconds and a minimum of 3.3 seconds. Thus, in the worst case (i.e., all sustained VT), the total computational time may have achieved up to 1505 h of computational time. By using a physiological ionic model, such as the TenTusccher-Panfilov model [67] or the ToR-ORd model [68], both available in CardioMat, the computational effort becomes about ten and twenty times higher, respectively, making the study unfeasible. Finally, virtual ablations were assumed to generate spherical lesions; however, lesions generated by catheter ablation may present more complex shapes depending on the modality and parameters adopted [47].

## 5. Conclusion

In this manuscript, we presented an image-based computational framework for conducting controlled in silico trials of ablation strategies in scar-related ventricular tachycardia. Starting from clinical imaging data, the proposed pipeline enables the generation of patient-specific electrophysiological substrates, the interactive implementation of virtual ablation lesions, and the standardized assessment of post-ablation VT reinducibility. Therefore, the proposed approach and software platform can be used to compare existing clinical strategies, test emerging substrate-based approaches, and quantitatively assess ablation efficacy together with ablation burden. As a proof of concept, we applied the framework to a cohort of patients with ischemic and non-ischemic cardiomyopathy and compared four substrate-based ablation strategies: scar homogenization, primary DZ ablation, primary and secondary DZ ablation, and CMR-guided scar dechanneling. The results demonstrated that all tested strategies reduced VT inducibility compared with baseline. Scar homogenization achieved the largest reduction in inducible VTs, but required the largest ablated volume. Conversely, CMR-guided scar dechanneling showed the most favorable efficiency profile, substantially reducing VT inducibility while limiting the amount of ablated viable myocardium. These findings highlight the importance of evaluating ablation strategies not only in terms of residual inducibility, but also in relation to the extent of myocardial tissue destruction. This study should not be interpreted as a definitive clinical ranking of substrate-based ablation strategies. Rather, it demonstrates that clinical imaging can be converted into patient-specific computational substrates suitable for controlled in silico trials. Within this framework, different ablation strategies can be implemented, visually inspected, quantitatively compared, and analyzed in terms of both post-ablation inducibility and ablation burden. The comparison among scar homogenization, DZ-based ablation, and CMR-guided scar dechanneling illustrates the type of mechanistic and quantitative information that can be extracted from the proposed platform. Future studies on larger virtual cohorts, ideally combined with prospective clinical validation, will be necessary to further assess the translational value of this approach and its potential role in image-guided VT ablation planning.

## Acknowledgments

This work was supported by the Italian Ministry of University and Research (MUR) in the framework of the FoReLab project (Departments of Excellence), the Regione Toscana under the program FSE+ Toscana 2021- 2027 (SICARDIO project) and the program PRS 2016-2020 (Bando ricerca salute 2018), and the European Union by the Next Generation EU project ECS00000017 ‘Ecosistema dell’Innovazione’ Tuscany Health Ecosystem (THE, PNRR, Spoke 9: Robotics and Automation for Health).

