## Supplementary Material for "An image-based framework for in silico trials of ablation strategies in scar-related ventricular tachycardia"

### Cardiac electrophysiology modeling

In this work, cardiac electrophysiology in the left ventricular models was simulated with the monodomain equation:

$$\frac{\partial V_M}{\partial t} - \nabla \cdot (D \nabla V_M) = -I_{ion}$$

where $V_M$ is the membrane potential, $D$ is the diffusion tensor, and $I_{ion}$ the ionic current.

The ionic current was represented by the phenomenological model we previously presented in [1, 2]. This phenomenological model has three state variables, including the transmembrane voltage, and its temporal evolution is described by three main currents: excitation, recovery, and transient outward. For the reader's convenience, we report below the full single-cell model equations.

$$\begin{cases} \dfrac{dV_M}{dt} = -I_{ion} + I_{stim} \\ \dfrac{du}{dt} = ke_{(V_M,u,w)}\left(\dfrac{V_M - B}{A_{(u)}} - u\right) \\ \dfrac{dw}{dt} = ke_{w(u)}\left(\dfrac{V_M - B}{A_{(u)}} - d_{w(u)}w\right) \end{cases}$$

The ionic currents are described by the following equations:

$$I_{ion} = g_{(u)}k(I_e + I_R + I_{To});$$

$$I_e = c_1(V_M - B)\left(a - \frac{V_M - B}{A_{(u)}}\right)\left(1 - \frac{V_M - B}{A_{(u)}}\right);$$

$$I_R = c_2u(V_M - B);$$

$$I_{To} = c_3w(V_M - B)s_{u(u)};$$

The model functions are described by the following equations:

$$A_{(u)} = A_0 + A_1u^2;$$

$$g_{(u)} = (\gamma_0 + \gamma_1 u)\frac{1 - \tanh\big(\alpha(u - \theta_u)\big)}{2} + g_0;$$

$$s_{u(u)} = \left(\frac{(u_M - u)}{u_M}\right)^2;$$

$$d_{w(u)} = d_w^0 / s_{u(u)};$$

$$e_w = g_{(u)} e_w^0;$$

$$e_{(V_M,u,w)} = \begin{cases} g_{(u)}(e_{11} + e_{12}|u|) \quad if \;\; du/dt \geq 0; dv/dt \geq 0; dw/dt \geq 0 \\ (e_{11} + e_{12}|u|) \; if \;\; du/dt \geq 0; dv/dt < 0; dw/dt < 0 \\ (e_{21} + e_{22}|u|) \;\; if \;\; du/dt \leq 0 \end{cases}$$

Local modifications of the parameters were introduced based on the normalized pixel signal intensity to replicate electrophysiological alterations observed in the border zone. In particular, the parameters $e_{11}^0$, $e_{12}^0$, $e_{21}$, $e_{22}$, $e_w^0$, $\gamma_0$, $\gamma_1$, $A_0$, $A_1$ were all multiplied by the corrective factor $(1 - NPI/4)$.

The baseline model parameters used for endocardial and epicardial tissue are reported in the table below:

| Parameter | Endocardium | Epicardium |
|---|---|---|
| $k$ | $1\ ms^{-1}$ | $1\ ms^{-1}$ |
| $c_1$ | 2.6 | 2.6 |
| $c_2$ | 1 | 1 |
| $c_3$ | 0.5 | 0.5 |
| $a$ | 0.18 | 0.18 |
| $A_0$ | 125 mV | 135 mV |
| $A_1$ | 60 mV | 0 mV |
| $B$ | -85 mV | -85 mV |
| $e_{11}^0$ | 0.001 | 0.0059 |
| $e_{12}^0$ | 0.0133 | 0 |
| $e_{21}$ | 0.0009 | 0.015 |
| $e_{22}$ | 0.0331 | 0 |
| $\gamma_0$ | 9.25 | 3 |
| $\gamma_1$ | 23.125 | 20 |
| $\alpha$ | 15 | 15 |
| $\theta_u$ | 0.2 | 0.2 |
| $g_0$ | 0.1 | 0.1 |
| $u_M$ | 0.58 | 0.58 |
| $e_w^0$ | 0.025 | 0.04 |
| $d_w^0$ | 5 | 0.6 |

### Preliminary analysis of fibrosis density effect

We conducted a preliminary analysis on 7 subjects to assess the effect of different fibrosis densities in the border zone on scar -related arrhythmogenesis. For each patient, we generated five computational models with different percentages of fibrotic tissue in the border zone. The density of border zone fibrosis was modulated according to the PSI, increasing close to the core scar. The fibrosis tissue percentage ranges used were 0-0% (i.e., without fibrosis), 0-20%, 20-40%, 40-60%, and 60-80%. Arrhythmia vulnerability was assessed for each model using a standardized protocol (S1 600 ms x 6, S2 350 ms, S3 300 ms, S4 280 ms) pacing from all 17 segments of the left ventricle. All the 595 3D simulations were set up and carried out with CardioMat [3]. A simulation was considered an arrhythmia if the reentry induced by the pacing protocol was maintained for 4 seconds.

A total of 72 sustained ventricular arrhythmias were induced, 49 of which were obtained with average fibrosis density equal to 50%. The number of events decreased very fast when border zone fibrosis density was reduced. Without considering fibrosis in the border zone, only one arrhythmic event was induced, whereas 4 were obtained in a single patient with average fibrosis density equal to 10%. Similarly, by increasing the fibrosis tissue percentages in the range 60-80% the number of events reduced to 7. The number of VT induced is significantly higher for border zone fibrosis percentage in the range 40-60% ($p<0.001$).

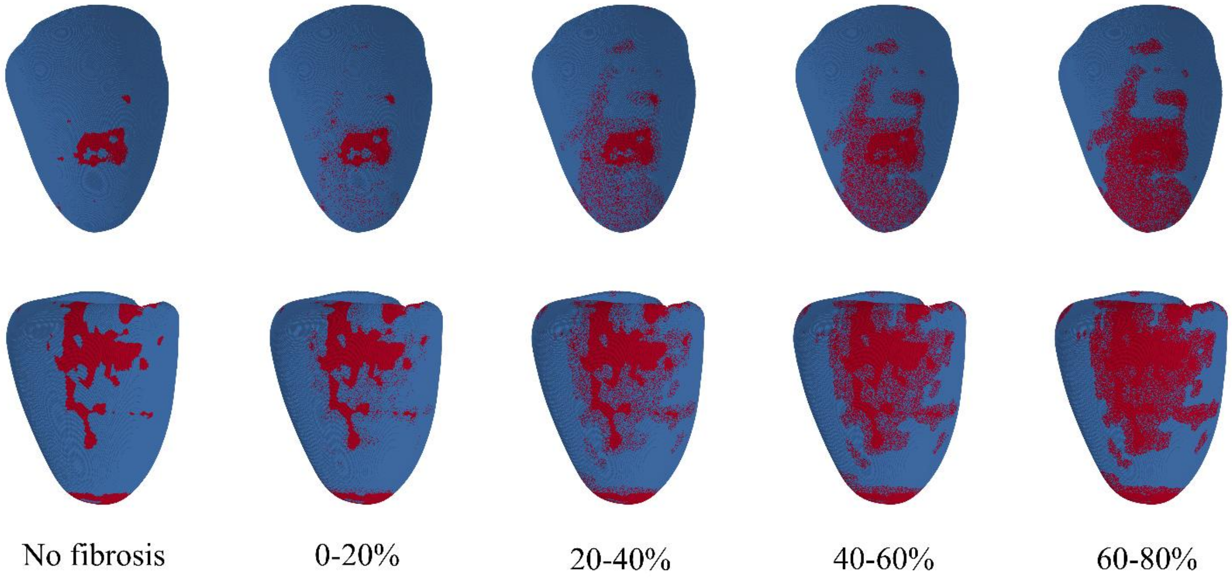


*Figure 1. Two patient models with different fibrosis density range in the border zone*

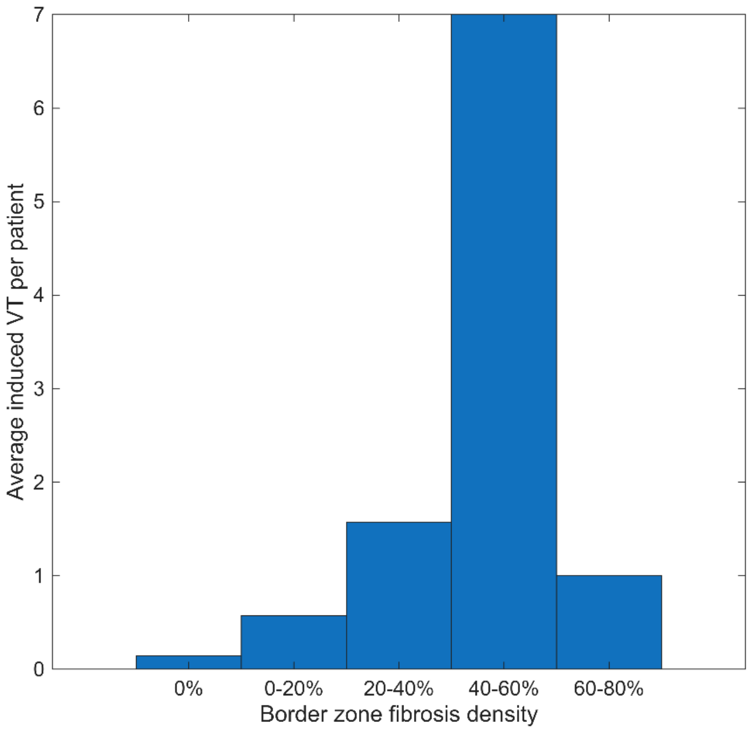


*Figure 2. Induced VT per patient for different border zone fibrosis density ranges*

### Complete patient-level results

The results of pre- and post-ablation VT inducibility testing are reported in the table below. For each patient, the number of sustained VTs and unique sustained VTs are reported prior and after each ablation strategy. The ablated volume is shown for each ablation, both in absolute and relative terms. We also introduced the number of ablated VTs per ml of ablated volume as a measure of ablation efficiency.

| ID | ISCH | NO ABLATION | | SCAR HOMOGENIZATION | | | | | | | |
|---|---|---|---|---|---|---|---|---|---|---|---|
| | | SUST. VT | UNIQUE VT | SUST. VT | UNIQUE VT | ABL VOL | REL. VOL | DELTA VT | DELTA VT % | ABL. VT PER ML | % VT / % ABL |
| 1 | 2 | 4 | 3 | 2 | 2 | 13,141 | 16 | 1 | 33,333 | 0,0761 | 0,0208 |
| 2 | 2 | 3 | 3 | 0 | 0 | 9,081 | 11,3 | 3 | 100 | 0,3304 | 0,0885 |
| 3 | 1 | 6 | 6 | 0 | 0 | 24,476 | 19,4 | 6 | 100 | 0,2451 | 0,0515 |
| 4 | 1 | 4 | 3 | 0 | 0 | 9,903 | 10,7 | 3 | 100 | 0,3029 | 0,0935 |
| 5 | 2 | 6 | 5 | 0 | 0 | 16,585 | 17,5 | 5 | 100 | 0,3015 | 0,0571 |
| 6 | 1 | 6 | 5 | 3 | 3 | 15,494 | 13,6 | 2 | 40 | 0,1291 | 0,0294 |
| 7 | 2 | 1 | 1 | 0 | 0 | 9,087 | 11,6 | 1 | 100 | 0,11 | 0,0862 |
| 8 | 2 | 0 | 0 | | | | | | | | |
| 9 | 2 | 10 | 9 | 1 | 1 | 21,25 | 20,3 | 8 | 88,889 | 0,3765 | 0,0438 |
| 10 | 1 | 8 | 3 | 0 | 0 | 12,725 | 14,8 | 3 | 100 | 0,2358 | 0,0676 |
| 11 | 2 | 10 | 6 | 2 | 1 | 19,615 | 25,1 | 5 | 83,333 | 0,2549 | 0,0332 |
| 12 | 1 | 0 | 0 | | | | | | | | |
| 13 | 1 | 0 | 0 | | | | | | | | |
| 14 | 1 | 13 | 12 | 9 | 6 | 27,018 | 14,5 | 6 | 50 | 0,2221 | 0,0345 |
| 15 | 1 | 11 | 7 | 2 | 1 | 30,568 | 25,5 | 6 | 85,714 | 0,1963 | 0,0336 |
| 16 | 2 | 14 | 4 | 0 | 0 | 21,091 | 26,97 | 4 | 100 | 0,1897 | 0,0371 |
| 17 | 1 | 9 | 8 | 7 | 4 | 37,094 | 36,74 | 4 | 50 | 0,1078 | 0,0136 |
| 18 | 2 | 12 | 8 | 0 | 0 | 37,525 | 37,56 | 8 | 100 | 0,2132 | 0,0266 |
| 19 | 1 | 6 | 2 | 0 | 0 | 7,153 | 16 | 2 | 100 | 0,2796 | 0,0625 |
| 20 | 2 | 4 | 3 | 0 | 0 | 31,401 | 38,94 | 3 | 100 | 0,0955 | 0,0257 |

| ID | DZ ABLATION (FIRST ROUND) | | | | | | | | DZ ABLATION (SECOND ROUND) | | | | | | | |
|---|---|---|---|---|---|---|---|---|---|---|---|---|---|---|---|---|
| | SUST. VT | UNIQUE VT | ABL VOL | REL. VOL | DELTA VT | DELTA VT % | ABL. VT PER ML | % VT / % ABL | SUST. VT | UNIQUE VT | ABL VOL | REL. VOL | DELTA VT | DELTA VT % | ABL. VT PER ML | % VT / % ABL |
| 1 | 7 | 3 | 5,064 | 6,2 | 0 | 0 | 0 | 0 | 5 | 2 | 8,11 | 9,8 | 1 | 33,333 | 0,1233 | 0,034 |
| 2 | 2 | 1 | 4,435 | 5,5 | 2 | 66,667 | 0,451 | 0,1212 | 2 | 2 | 5,384 | 6,7 | 1 | 33,333 | 0,1857 | 0,0498 |
| 3 | 4 | 3 | 7,428 | 5,9 | 3 | 50 | 0,4039 | 0,0847 | 5 | 3 | 9,248 | 7,3 | 3 | 50 | 0,3244 | 0,0685 |
| 4 | 0 | 0 | 7,108 | 7,7 | 3 | 100 | 0,4221 | 0,1299 | 0 | 0 | 7,108 | 8 | 3 | 100 | 0,4221 | 0,125 |
| 5 | 5 | 3 | 5,213 | 5,5 | 2 | 40 | 0,3837 | 0,0727 | 7 | 5 | 7,313 | 7,7 | 0 | 0 | 0 | 0 |
| 6 | 9 | 5 | 7,431 | 6,5 | 0 | 0 | | 0 | 8 | 3 | 9,45 | 8,3 | 2 | 40 | 0,2116 | 0,0482 |
| 7 | 0 | 0 | 7,759 | 9,9 | 1 | 100 | 0,1289 | 0,101 | 0 | 0 | 7,759 | 9,9 | 1 | 100 | 0,1289 | 0,101 |
| 8 | | | | | | | | | | | | | | | | |
| 9 | 10 | 8 | 3,6 | 3,4 | 1 | 11,111 | 0,2778 | 0,0327 | 10 | 5 | 11,663 | 11,1 | 4 | 44,444 | 0,343 | 0,04 |
| 10 | 2 | 2 | 5,085 | 5,9 | 1 | 33,333 | 0,1967 | 0,0565 | 0 | 0 | 9,858 | 11,5 | 3 | 100 | 0,3043 | 0,087 |
| 11 | 11 | 4 | 4,457 | 5,7 | 2 | 33,333 | 0,4487 | 0,0585 | 10 | 5 | 7,004 | 9 | 1 | 16,667 | 0,1428 | 0,0185 |
| 12 | | | | | | | | | | | | | | | | |
| 13 | | | | | | | | | | | | | | | | |
| 14 | 12 | 9 | 14,5 | 7,8 | 3 | 25 | 0,2069 | 0,0321 | 10 | 8 | 24,077 | 12,9 | 4 | 33,333 | 0,1661 | 0,0258 |
| 15 | 10 | 4 | 6,112 | 5,1 | 3 | 42,857 | 0,4908 | 0,084 | 8 | 2 | 20,409 | 17 | 5 | 71,429 | 0,245 | 0,042 |
| 16 | 9 | 1 | 8,353 | 10,68 | 3 | 75 | 0,3592 | 0,0702 | 0 | 0 | 11,182 | 14,3 | 4 | 100 | 0,3577 | 0,0699 |
| 17 | 8 | 6 | 8,767 | 8,68 | 2 | 25 | 0,2281 | 0,0288 | 6 | 5 | 18,105 | 17,93 | 3 | 37,5 | 0,1657 | 0,0209 |
| 18 | 8 | 5 | 13,709 | 13,72 | 3 | 37,5 | 0,2188 | 0,0273 | 7 | 5 | 15,019 | 15,03 | 3 | 37,5 | 0,1997 | 0,025 |
| 19 | 8 | 1 | 3,121 | 6,98 | 1 | 50 | 0,3204 | 0,0716 | 0 | 0 | 5,314 | 11,89 | 2 | 100 | 0,3764 | 0,0841 |
| 20 | 2 | 1 | 13,399 | 16,62 | 2 | 66,667 | 0,1493 | 0,0401 | 0 | 0 | 16,356 | 20,28 | 3 | 100 | 0,1834 | 0,0493 |

| ID | DECHANNELING ABLATION | | | | | | | |
|---|---|---|---|---|---|---|---|---|
| | SUST. VT | UNIQUE VT | ABL VOL | REL. VOL | DELTA VT | DELTA VT % | ABL. VT PER ML | % VT / % ABL |
| 1 | 3 | 2 | 5,525 | 6,7 | 1 | 33,333 | 0,181 | 0,0498 |
| 2 | 2 | 1 | 3,839 | 4,2 | 2 | 66,667 | 0,521 | 0,1587 |
| 3 | 6 | 2 | 8,772 | 7 | 4 | 66,667 | 0,456 | 0,0952 |
| 4 | 0 | 0 | 6,403 | 6,9 | 3 | 100 | 0,4685 | 0,1449 |
| 5 | 3 | 2 | 4,001 | 4,2 | 3 | 60 | 0,7498 | 0,1429 |
| 6 | 1 | 1 | 12,67 | 11,2 | 4 | 80 | 0,3157 | 0,0714 |
| 7 | 0 | 0 | 6,792 | 8,6 | 1 | 100 | 0,1472 | 0,1163 |
| 8 | | | | | | | | |
| 9 | 5 | 3 | 17,074 | 16,3 | 6 | 66,667 | 0,3514 | 0,0409 |
| 10 | 2 | 1 | 7,068 | 8,2 | 2 | 66,667 | 0,283 | 0,0813 |
| 11 | 6 | 4 | 9,845 | 12,6 | 2 | 33,333 | 0,2031 | 0,0265 |
| 12 | | | | | | | | |
| 13 | | | | | | | | |
| 14 | 11 | 10 | 14,994 | 8 | 2 | 16,667 | 0,1334 | 0,0208 |
| 15 | 7 | 3 | 13,101 | 10,9 | 4 | 57,143 | 0,3053 | 0,0524 |
| 16 | 3 | 2 | 8,956 | 11,45 | 2 | 50 | 0,2233 | 0,0437 |
| 17 | 7 | 6 | 15,666 | 15,52 | 2 | 25 | 0,1277 | 0,0161 |
| 18 | 6 | 3 | 14,572 | 14,59 | 5 | 62,5 | 0,3431 | 0,0428 |
| 19 | 8 | 1 | 2,691 | 6,02 | 1 | 50 | 0,3716 | 0,0831 |
| 20 | 6 | 2 | 11,049 | 13,7 | 1 | 33,333 | 0,0905 | 0,0243 |

**Complete patient-level results.** Each patient corresponds to a row beginning with the patient ID number. The second column specifies if the patient's scar is of ischemic (1) or non ischemic (2) nature. The subsequent columns report the results of the simulations conducted in our in silico trial study. The following abbreviations have been used: SUST. VT: sustained VT; UNIQUE VT: unique sustained VT; ABL VOL: ablation volume in ml REL VOL: relative ablation volume (normalized to the total left ventricular volume); DELTA VT: ablated VT; DELTA VT %: percentage of ablated VT normalized to the total number of VTs induced; ABL VT PER ML: number of ablated VTs normalized to ml of ablated volume;%VT/%ABL: percentage of ablated VTs normalized to the relative ablated volume.

### Computational accounting

In silico VT inducibility testing prior to ablation was performed on all the models generated from imaging data of the 20 enrolled patients. 3 patients were not inducible. Thus, in silico virtual ablation was performed only on the 17 inducible patients following three different strategies: scar homogenization, cardiac magnetic resonance-guided dechanneling, and primary deceleration zones ablation. Patients who showed residual VT inducibility after primary deceleration zone ablation (15 patients) also underwent virtual ablation of secondary deceleration zones. Thus, in total, we conducted 86 in silico VT inducibility tests. Each VT inducibility test accounts for 17 simulations, delivering the stimulation protocol in a different AHA segment. Therefore, we carried out 1462 simulations, 414 resulting in sustained VT. Each simulation can last up to 8.33 seconds of simulated time in case of sustained VT. In case no arrhythmia was induced, a single simulation covered 3.33 seconds. 1s of simulation requires, on average, about 450 seconds to be completed on a NVIDIA GeForce 3090 GPU. Thus, only the 414 sustained VT simulations required approximately 431computational hours. A similar amount was requested for the remaining simulations, leading to a total computational account of approximately 800 computational hours.

### Supplementary Video

The manuscript is associated with three supplementary videos:

- Video S1 shows the use of the AblateTool standalone application
- Video S2 shows the example presented in Figure 11 of the manuscript
- Video S3 shows an example of fibrillation pattern (the same as Figure 7 of the manuscript)

[1] N. Biasi and A. Tognetti, "A computationally efficient dynamic model of human epicardial tissue," *Plos one,* vol. 16, no. 10, p. e0259066, 2021.

[2] N. Biasi, P. Seghetti, M. Mercati, and A. Tognetti, "A reaction-diffusion heart model for the closed-loop evaluation of heart-pacemaker interaction," *IEEE Access,* vol. 10, pp. 121249-121260, 2022.

[3] N. Biasi, P. Seghetti, M. Parollo, G. Zucchelli, and A. Tognetti, "A Matlab Toolbox for cardiac electrophysiology simulations on patient-specific geometries," *Computers in Biology and Medicine,* vol. 185, p. 109529, 2025.
